# Jasmine: Population-scale structural variant comparison and analysis

**DOI:** 10.1101/2021.05.27.445886

**Authors:** Melanie Kirsche, Gautam Prabhu, Rachel Sherman, Bohan Ni, Sergey Aganezov, Michael C. Schatz

## Abstract

The increasing availability of long-reads is revolutionizing studies of structural variants (SVs). However, because SVs vary across individuals and are discovered through imprecise read technologies and methods, they can be difficult to compare. Addressing this, we present Jasmine (https://github.com/mkirsche/Jasmine), a fast and accurate method for SV refinement, comparison, and population analysis. Using an SV proximity graph, Jasmine outperforms five widely-used comparison methods, including reducing the rate of Mendelian discordance in trio datasets by more than five-fold, and reveals a set of high confidence *de novo* SVs confirmed by multiple long-read technologies. We also present a harmonized callset of 205,192 SVs from 31 samples of diverse ancestry sequenced with long reads. We genotype these SVs in 444 short read samples from the 1000 Genomes Project with both DNA and RNA sequencing data and assess their widespread impact on gene expression, including within several medically relevant genes.

## Introduction

Structural variants (SVs) are defined as large-scale genomic mutations affecting more than 30 to 50 basepairs, and include insertions, deletions, duplications, inversions, and translocations (Alonge et al. 2020; Alkan, Coe, and Eichler 2011). Such variants are responsible for more divergent basepairs across human genomes than any other class of variation (Chiang et al. 2017), and have been associated with many major diseases and phenotypes, including cancer (Aganezov et al. 2020; Nattestad et al. 2018) and autism (Brandler et al. 2018). They have also been shown to have phenotypic effects in other species, such as increased fruit size in tomato (Alonge et al. 2020) or altered growth under stress in yeast (Jeffares et al. 2017). However, much of the impact of structural variants remains unknown because of the inability of SVs in complex regions to be accurately identified by short reads, which make up the majority of existing genomic sequencing data (Sedlazeck, Lee, et al. 2018; Mahmoud et al. 2019).

In recent years, the emergence of long-read genomic sequencing technologies (Korlach et al. 2010; M. Jain et al. 2016; Wenger et al. 2019; Goodwin, McPherson, and McCombie 2016) and the development of specialized software for alignment (C. Jain et al. 2020; Sedlazeck, Rescheneder, et al. 2018; Li 2018) and variant calling (Sedlazeck, Rescheneder, et al. 2018; Jiang et al. 2020) have enabled the characterization of complex structural variants which were difficult or impossible to study from short reads alone (Sedlazeck, Lee, et al. 2018). For this reason, many population variant inference studies include long-read sequencing data for multiple individuals instead of or in addition to short-read data (Chaisson et al. 2019; Audano et al. 2019; Beyter et al. 2021).

Because there are multiple sequencing technologies, aligners, and SV callers that could be used, SV-processing pipelines for population-scale studies are frequently optimized for the particular dataset being analyzed (Jeffares et al. 2017; Beyter et al. 2021), making it difficult to compare SVs called in different studies or to accurately screen newly sequenced samples for known variants. In addition, existing tools for comparing SV callsets from different samples have issues such as collapsing multiple variants in the same individual, including variants of different types, and producing callsets that vary substantially when the order of the input samples is changed. As the cost of long-read sequencing continues to fall and the number of population-scale SV studies continues to rise, there is an increasingly apparent need for methods which can accurately compare variants across a range of datasets.

To address this need, we introduce an optimized software pipeline for accurately detecting SVs and comparing these variant calls across large numbers of individuals (**Figure 1**). This pipeline enhances existing methods for alignment (C. Jain et al. 2020) and variant calling (Sedlazeck, Rescheneder, et al. 2018) with new methods for refining the sequences and breakpoints of SV calls, and for comparing variant calls between different individuals to achieve a unified callset. The first new method, Iris, refines variant calls by gathering the set of reads that support each variant’s presence and using them to polish the variant sequence. The second major novel method, Jasmine, compares and merges calls in different individuals corresponding to the same variant. Jasmine improves upon other SV merging methods by representing variants as points in space based on their breakpoints and lengths and constructing a graph of SV proximity, where edges represent pairs of SVs with a small Euclidean distance between them. To avoid the high time and memory overhead of computing and storing the entire graph, Jasmine uses a KD-Tree (Bentley 1975) to dynamically locate nearby variant pairs and implicitly detect low-weight edges. Jasmine then treats the comparison/merging problem as one of finding a minimal-weight acyclic subgraph of the proximity graph which satisfies certain constraints, and solves this problem with a constrained version of Kruskal’s algorithm for minimum spanning trees (Kruskal 1956). Both Iris and Jasmine are available as stand-alone software packages and are available within bioconda.

**Figure 1:**
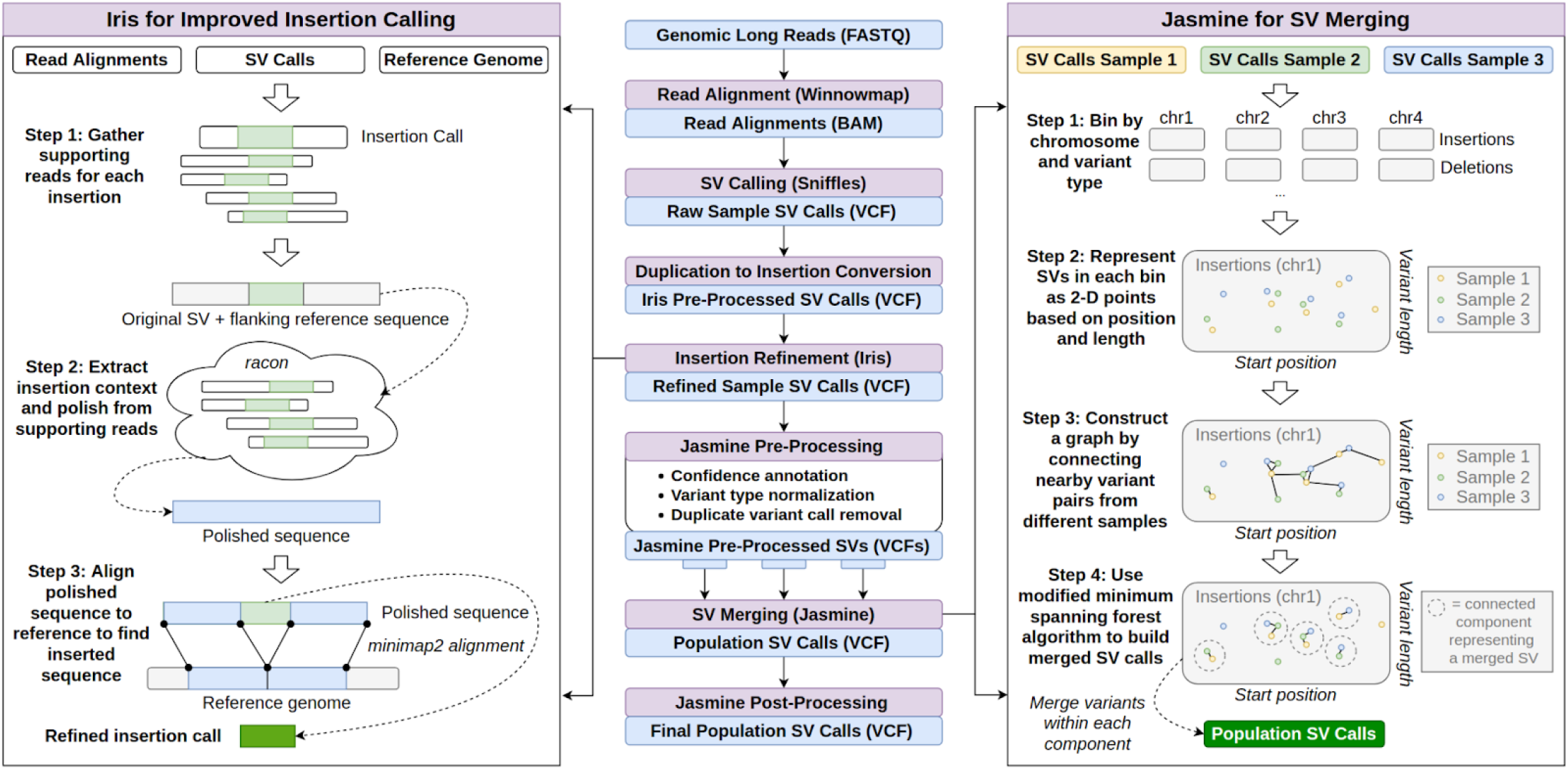
SV Inference Pipeline. This pipeline produces population-level SV calls from FASTQ files using a number of existing methods as well as two novel methods, Iris and Jasmine. Iris uses consensus methods to improve the accuracy of the breakpoints and sequence of insertion SVs. Jasmine uses a graph of SV proximity and a constrained minimum spanning forest algorithm to compare and combine variants across multiple individuals.

Using a combination of simulated and real datasets, we show that this pipeline produces more accurate SV calls than several widely used methods across a variety of metrics. First, by applying our methods to a HiFi dataset from the HG002 Genome-In-A-Bottle (GIAB) Ashkenazim trio, we illustrate that our approach achieves a five-fold reduction in the number of Mendelian discordant variants, while identifying multiple high-confidence *de novo* variants in the child supported by three independent sequencing platforms. We also analyze this trio to identify signatures of variants specifically derived from each particular technology. This enables us to establish recommended variant calling parameters for different sequencing technologies which minimize Mendelian discordance as well as false merges.

We next show that Jasmine improves SV merging and addresses the major issues that other methods encounter when scaling up to large cohorts. We call variants with our pipeline from publicly available long-read data for 31 samples, and generate a panel of long-read SV calls which can be used for screening further samples. Finally, we genotype this SV panel in 444 high coverage short-read samples from the 1000 Genomes Project (Byrska-Bishop et al. 2021) and discover thousands of novel SV associations with gene expression. Many of these SVs have CAVIAR posterior probabilities of causality that exceed those of previously reported SNPs, indicating likely functional relevance. This includes a deletion associated with the expression of SEMA5A, which has been implicated as an autism susceptibility gene (Melin et al. 2006), as well as within several other genes of interest.

## Results

### Reduced Mendelian Discordance in an Ashkenazim Trio

A common application of SV and other variant inference methods is the identification of *de novo* variants, or variants which are present in an individual but neither of their parents. Such variants have been associated with autism (Iossifov et al. 2014) and cancer (Renaux-Petel et al. 2018), and *de novo* variant analysis is frequently used as a starting point for identifying the cause of genetic diseases or other phenotypes of interest (Veltman and Brunner 2012). However, because of shortcomings in SV inference and comparison methods, identifying *de novo* SVs remains a difficult problem. For example, one widely used pipeline consisting of ngmlr, sniffles (Sedlazeck, Rescheneder, et al. 2018), and SURVIVOR (Jeffares et al. 2017) gives thousands of candidate *de novo* variants when applied to high-accuracy HiFi sequencing data from the HG002 Ashkenazim trio (**Figure 2a**). Because the number of *de novo* SVs is typically estimated to be less than ten per generation on average (Belyeu et al. 2021), almost all of these variant calls are either false positives in the child, false negatives in one or both parents, or errors in merging the callsets. Collectively, we refer to these false outcomes as Mendelian discordant variants.

**Figure 2.**
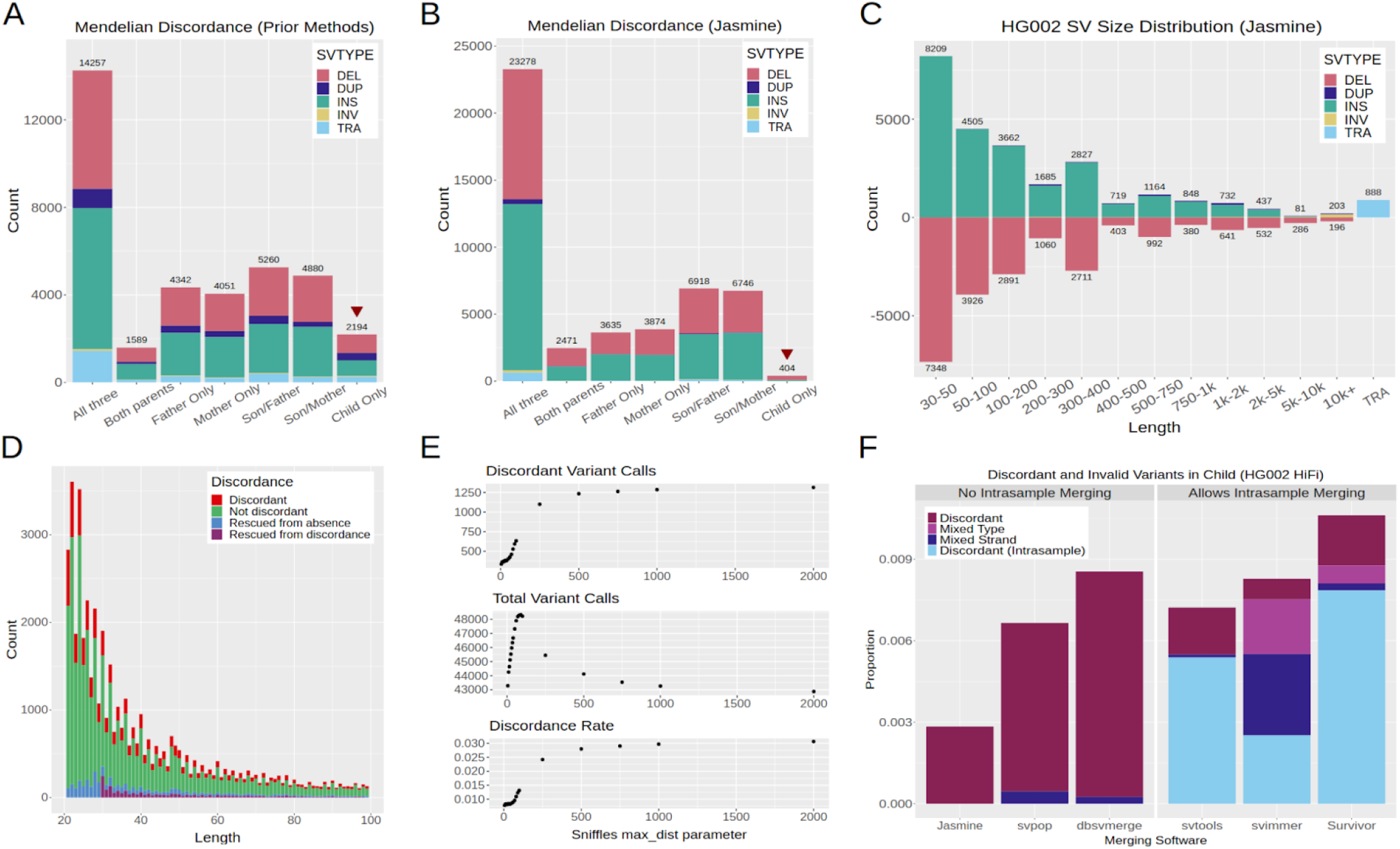
Mendelian Discordance in the HG002 Ashkenazim Trio. We called SVs from HiFi data for the Ashkenazim trio consisting of HG002 (son - 46,XY), HG003 (father - 46,XY), and HG004 (mother - 46,XX) using several prior methods as well as our pipeline. **a.)** The number of samples called in each subset of individuals when using prior methods: ngmlr for alignment, sniffles for SV calling, and SURVIVOR for consolidating SVs between samples. **b.)** The number of samples called in each subset of individuals when using our optimized pipeline. **c.)** The distribution of SV types and lengths in HG002 with our pipeline. **d.)** The benefits of using “double thresholding” to improve variant discovery in HG002 while also reducing the rate of Mendelian discordance. “Rescued from absence” refers to SVs which would have been missed in HG002 using a single threshold. “Rescued from discordance” refers to SVs which would have been discordant in HG002 with a single threshold, but which we were able to detect in one or both parents with double thresholding. **e.)** The effects of the sniffles *max_dist* parameter on downstream discordance. Using a tighter bound of 50 on the maximum distance sniffles allows between breakpoints in individual reads increases the total number of variants discovered while at the same time reducing the number of discordant variants compared to the default value of 1000. **f.)** The rate of discordance when comparing SVs between individuals with Jasmine as well as five existing methods for population inference. Jasmine reduces the discordance rate while at the same time addressing issues present in other methods such as merging variants of different types, variants with the same type but corresponding to unique breakpoint adjacencies (mixed strand), or variants within the same sample.

To address the large number of discordant variants, our optimized pipeline offers a number of improvements which reduce the rate of Mendelian discordance by more than a factor of five (**Figure 2b**) while discovering significantly more SVs (**Figure 2c**). These improvements include the mitigation of threshold effects (**Figure 2d**), improved variant calling parameters (**Figure 2e**), and using Jasmine for SV merging (**Figure 2f**). Furthermore, we compared Jasmine to five existing methods (Shi et al. 2021; Jeffares et al. 2017; Ebert et al. 2021; Larson et al. 2019; Beyter et al. 2021) for SV comparison between samples, and Jasmine achieves the lowest rate of discordance and correctly avoids merging variants of different types or variants from the same sample. In addition, Jasmine avoids merging variants of the same type which correspond to unique breakpoint adjacencies, which is particularly important when resolving complex nested SVs (**Supplementary Figure 21**). The resulting reduction in Mendelian discordant variants enables *de novo* variants to be identified more easily, as it is typically necessary to screen all discordant variants manually when searching for true *de novo* variants.

### SV Analysis Across Sequencing Technologies

Improved methods for comparing multiple SV callsets also enable the comparison of variants identified in a single individual from different sequencing technologies. We evaluated three different technologies applied to HG002: Pacific Biosciences Continuous Long Reads (CLR), Pacific Biosciences High-Fidelity (HiFi) circular consensus sequencing and Oxford Nanopore long reads (ONT) basecalled with Guppy 4.2.2. Variants were called separately from each technology, and the resulting callsets were merged with Jasmine. The three callsets were largely in agreement, with 30,590 out of 46,906 variants being supported by all three technologies (**Figure 3a and 3b**). The set of technology-concordant variants, shown in **Figure 3c**, shows that insertion and deletion calls are largely balanced, with a slight enrichment of insertions, shown in previous studies to be caused by missing sequence in the human reference genome (Audano et al. 2019). There is also an increased number of variants around sizes of 300bp and 6-7kbp, corresponding to SINE and LINE elements respectively.

**Figure 3.**
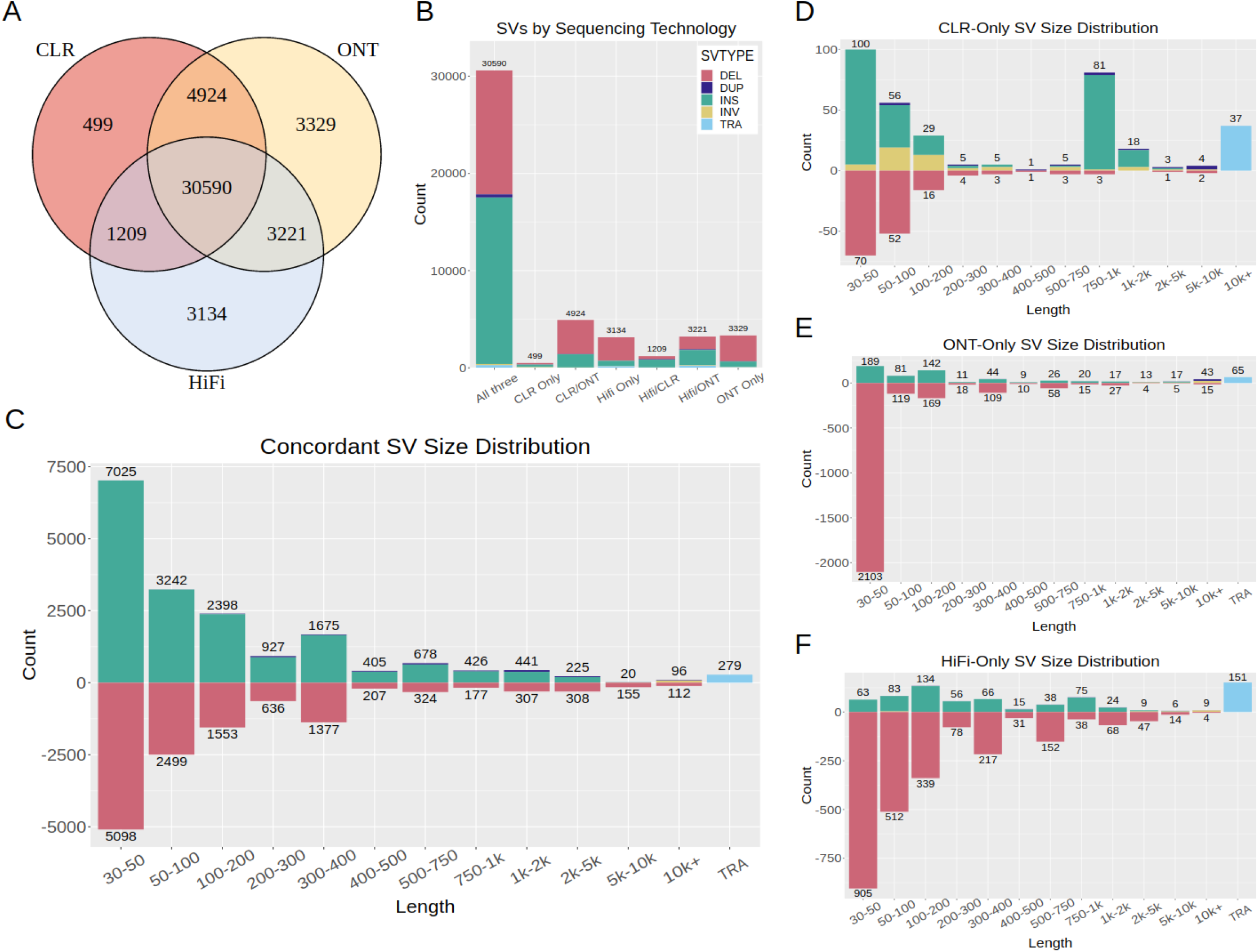
SV Inference across Sequencing Technologies in HG002. We called SVs in HG002 separately from Pacbio CLR data, Oxford Nanopore data, and Pacbio HiFi CCS data, and used Jasmine to compare the variants discovered by each of them. **a.)** The number of variants discovered by each subset of technologies. **b.)** The variant type distribution within each subset of technologies. **c.)** The distribution of types and lengths among SVs for which all of the technologies agree. **d-f.)** The SV type and length distributions for SVs unique to CLR, ONT, and HiFi respectively.

We also examined variants that were identified only by a single technology, as these may reveal systematic biases in variant calling caused by each technology’s error model. Of the 499 variants identified exclusively in CLR data (**Figure 3d**), there were 244 insertions and 155 deletions, with an excess of insertions in the size range 750 to 1000, corresponding to a known error characteristic of CLR sequencing (Sedlazeck, Rescheneder, et al. 2018). Of the 3329 ONT-only variant calls (**Figure 3e**), there were 539 insertions and 2652 deletions, with an enrichment of small deletions less than 50 basepairs in length. In addition, we found that many of the variants, particularly deletions, unique to ONT or HiFi are in centromeric regions or satellite repeats (**Supplementary Figure 13**).

### *De Novo* Variant Discovery

We next leveraged our methods, as well as data from all three technologies listed above, to screen the HG002 trio for *de novo* variants. We called variants from each of the three technologies in HG002 as well as both parents, for a total of nine callsets. We merged these nine callsets with Jasmine and filtered out any variants which were present in one or more of the six parent callsets. Of the remaining variants, we stratified them by which technologies supported their presence in the child and found that there were 16 which were supported by all three technologies (**Figure 4a**), with an additional 35 that were supported by HiFi and at least one other technology, a 43-fold reduction in candidates from using prior methods (**Figure 2a**).

**Figure 4.**
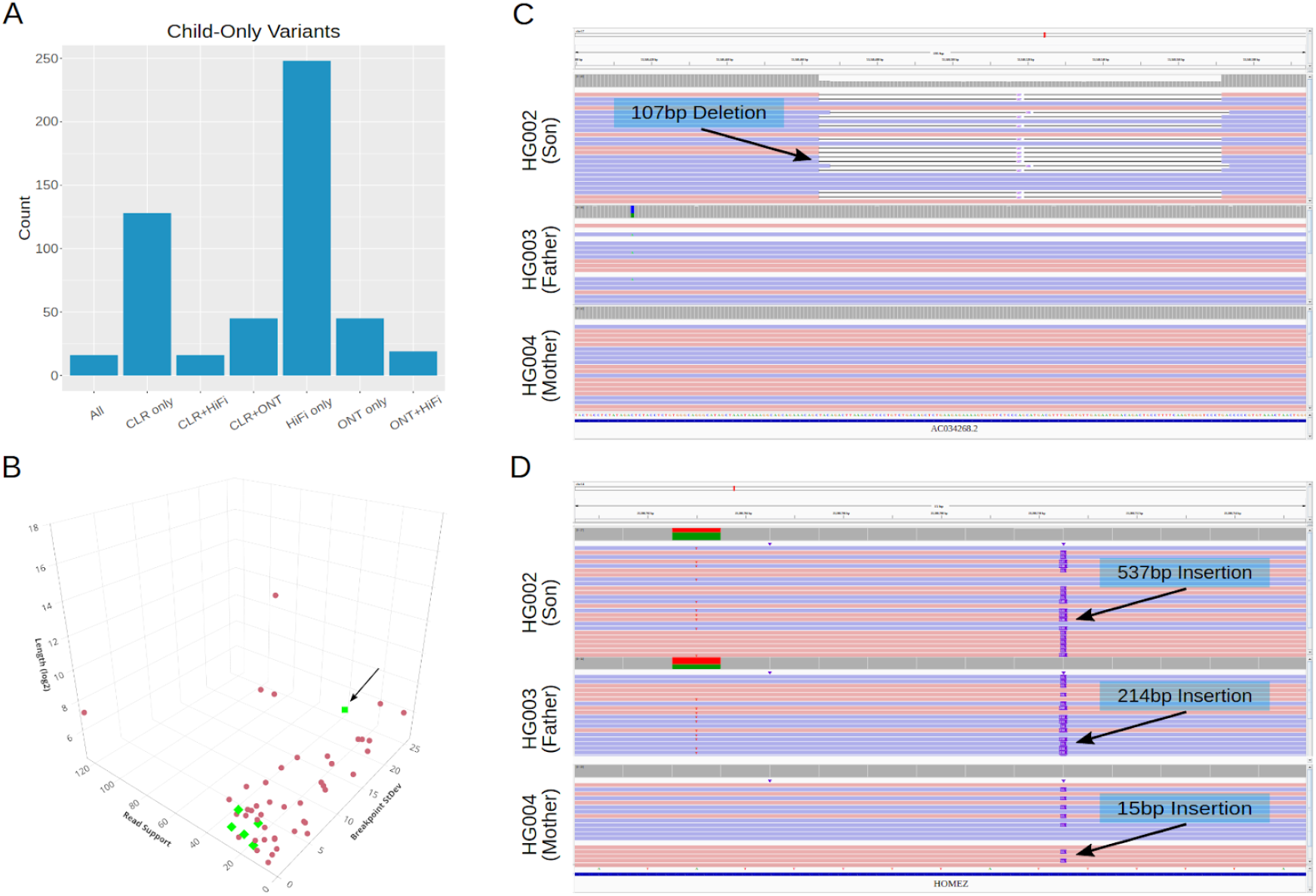
*De Novo* SV Discovery in HG002. We called SVs in each of HG002, HG003, and HG004 from three different sequencing technologies - CLR, ONT, and HiFi - to identify potential *de novo* variants that were called in none of the six parent callsets but one or more of the HG002 callsets. **a.)** The number of SVs which are absent in all six parent callsets whose presence in HG002 is supported by each subset of technologies. While we manually inspected all SVs supported by HiFi and at least one other technology, both of the examples in (a) and (b) were supported by all three technologies. **b.)** All SVs supported by HiFi and at least one other technology in HG002 that are absent in all parent callsets. The potential *de novo* SVs we identified are highlighted in green, with the microsatellite repeat expansion denoted by an arrow. **c.)** A potential *de novo* 107bp deletion in HG002 at chr17:53340465. **d.)** A potential *de novo* microsatellite repeat expansion in HG002 at chr14:23280711.

Upon manual inspection, six of these were high confidence *de novo* SVs (**Figure 4b**), while the remaining candidates were in noisy regions that displayed split read alignments, but we could not be certain whether the alignments were correct (**Supplementary Figure 16**). One of the high-confident candidates, a 107bp deletion at chr17:53340465 (**Figure 4c**), was previously identified as a *de novo* SV in a previous effort to characterize the variants in HG002 (Zook et al. 2020). Another example, a 537bp insertion at chr14:23280711, consists of a microsatellite repeat expansion on the paternal haplotype, a known class of mutations often caused by replication slippage (Ellegren 2004) (**Figure 4d**). These and other examples (**Supplementary Figures 14-16**) show that our approach can correctly identify known *de novo* SVs as well as identify potential *de novo* variants which were previously undiscovered.

### Population SV Inference

As the cost of long-read sequencing has continued to decrease in recent years, long-read studies including large cohorts have become more prevalent (Shi et al. 2021; Beyter et al. 2021). As this trend is expected to continue (Ranallo-Benavidez et al. 2021), it is particularly important for SV inference methods to be able to scale to many samples. To compare Jasmine to existing approaches, we called SVs in 31 publicly available long-read samples (**Supplementary Table 2**) and observed the results of merging these callsets with each method. All methods produced a population-level callset within a few hours with 24 threads on a modern 4GHz server with 192GB of RAM, but the callsets produced by existing approaches suffer from a number of issues. In addition to the invalid merges mentioned above (**Figure 2d**), several of the existing methods use algorithms which give different merging results, and consequently different numbers of total variant calls, based on the input order of the sample callsets (**Figure 5a**). This problem only worsens as the number of samples grows and the number of possible sample orderings increases exponentially. Conversely, Jasmine’s algorithm, which merges variant pairs in increasing order of their breakpoint distances irrespective of the input order, produces identical results after any permutation of input files. Finally, there is an abundance of low-confidence likely false positive SV calls in samples sequenced with CLR (**Supplementary Figure 17**), and methods which use a constant breakpoint distance threshold incorrectly merge these calls with high-confidence SV calls in other samples to obtain an unreasonable trimodal allele frequency distribution (**Supplementary Figure 18**).

**Figure 5.**
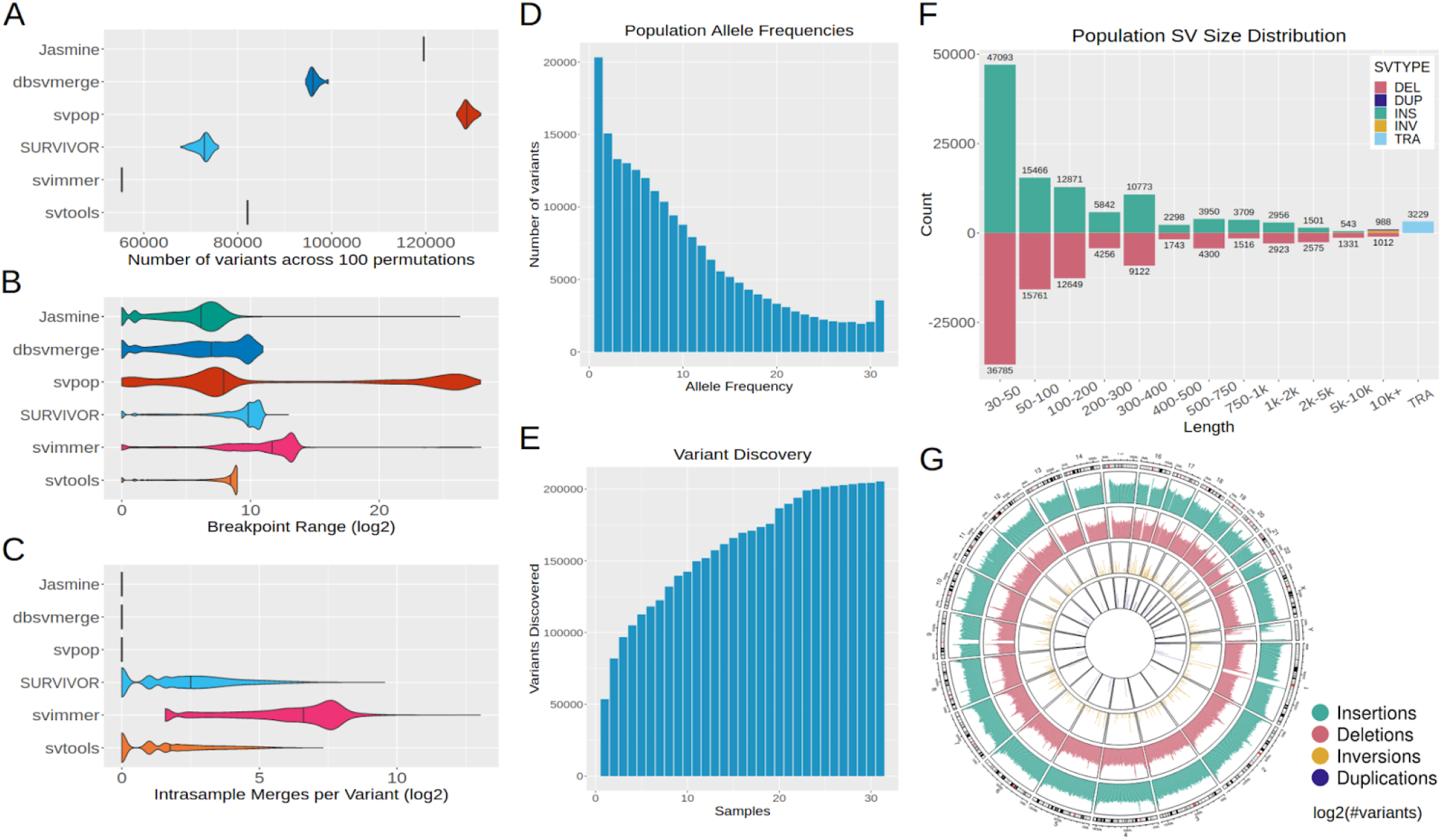
Population-Scale Inference from Public Datasets. We called SVs with our pipeline in a cohort of 31 samples from diverse ancestries and sequencing technologies and used Jasmine as well as five prior methods to combine the individual samples’ SVs into a population-scale callset. **a.)** The number of SVs obtained with each merging software across 100 runs with the list of input VCFs randomly shuffled each time. **b.)** The distribution of the range of breakpoints of SV calls merged into single variants by each software, excluding unmerged variants. **c.)** The number of intrasample merges within single merged variants, defined as the number of variants minus the number of unique samples, for each software. **d.)** The allele frequency distribution of variants merged by Jasmine. **e.)** The number of SVs discovered by Jasmine as the number of samples increases. **f.)** The distribution of SV types and lengths in the cohort when using Jasmine. **g.)** The number of SVs in the cohort in 1Mbp bins across the human genome.

Using our SV inference pipeline, we created a panel of long-read SVs from these 31 samples. The datasets we used include individuals from a wide range of ancestral backgrounds, as well as sequencing data from multiple technologies. Variants were called in each sample separately and merged with Jasmine to create a unified callset. The allele frequency distribution is monotonically decreasing as expected, except an excess of variants at 100% corresponding to errors and/or minor alleles in the reference (Audano et al. 2019) (**Figure 5d**). The cumulative number of variants increases with the number of samples, but at a decreasing rate (**Figure 5e**). The indels are approximately balanced, with a slight bias towards insertions, and there are spikes in the size distribution around 300bp and 6-7kbp for SINE and LINE elements (**Figure 5f**). There is also an enrichment of SVs in the centromeres and telomeres (**Figure 5g**), likely due to a combination of missing reference sequence, repetitive sequence which is difficult to align to, and greater recombination rates (Audano et al. 2019).

### Measuring Effects of SVs on Gene Expression

Previous expression quantitative trait loci (eQTL) studies have shown that SVs often have large effects on gene expression and that they are causal at 3.5-6.8% of eQTLs (Consortium and The 1000 Genomes Project Consortium 2015; Chiang et al. 2017). To investigate this with our enhanced catalog of SVs, we used Paragraph (Chen et al. 2019) to genotype each SV in 444 individuals from the 1000 Genomes Project (1KGP) for which gene expression data is publicly available (Lappalainen et al. 2013), after removing SVs that were inconsistent with population genetics expectations based on the Hardy-Weinberg equilibrium (**Figure 6a**). Following the prior studies, we mapped SV-eQTLs by pairing common (MAF ≥ 0.05) SVs to genes within 1 Mbp using gene expression data in lymphoblastic cell lines from the GEUVADIS consortium (Lappalainen et al. 2013). We then fit a linear model to measure the effect sizes of these SVs on gene expression, and found that 5,456 pairs passed a significance threshold with 10% FDR, which is substantially higher than the 478 pairs that we observe among short-read SVs. These associations occur for both deletions and insertions, and both have approximately the same effect size distribution (**Figure 6b**). These data suggest that many of the SVs that are only visible through genotyping long-read-based variant calls have large effects on gene expression and thus are potentially functionally relevant.

**Figure 6.**
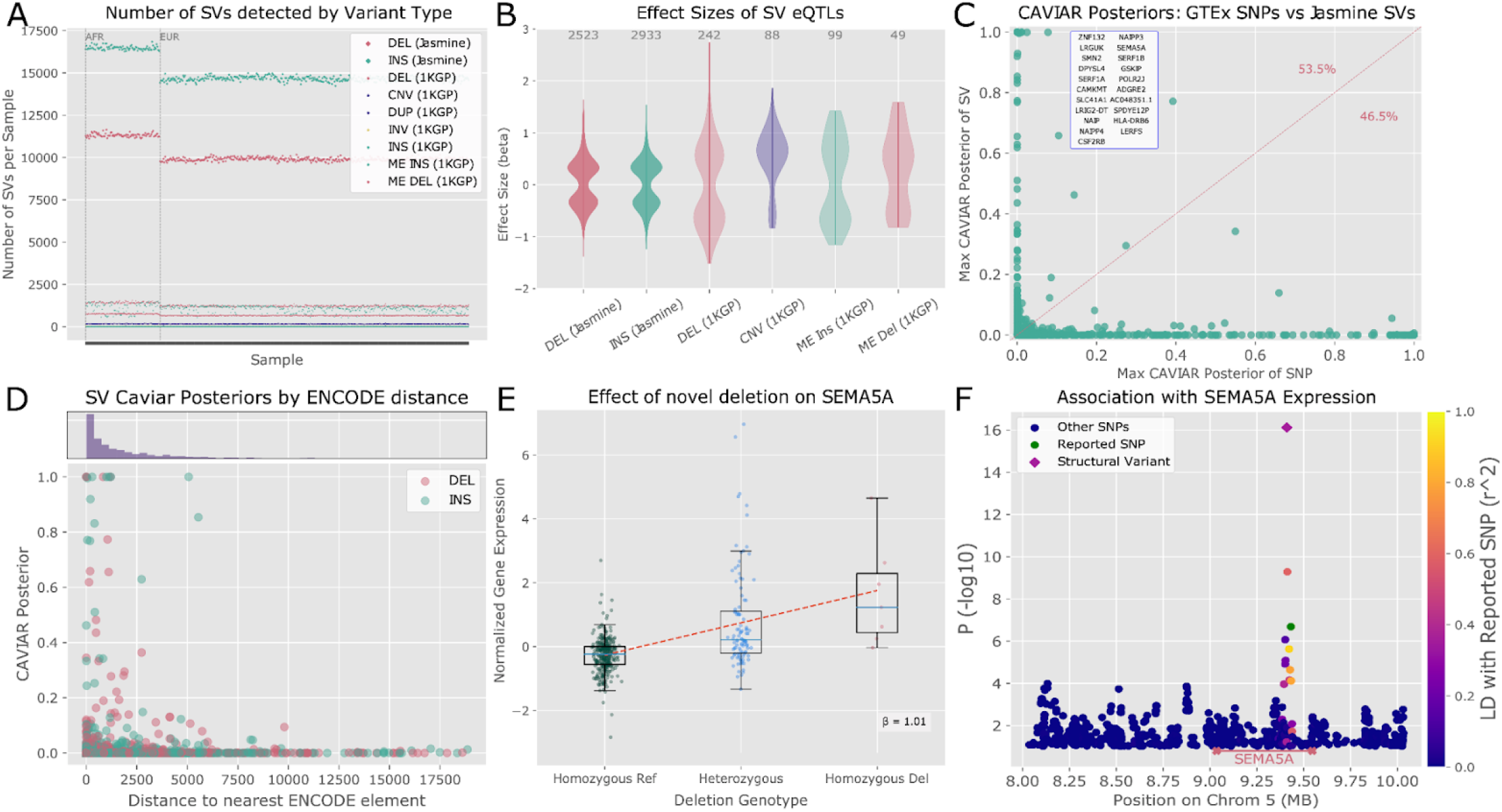
Functional impact of SVs from Jasmine. We used Paragraph to genotype SVs from the cohort of 31 samples in 444 samples from the 1000 Genomes Project which have RNA-seq data. **a.)** Number of SVs detected per sample for genotyped SVs (Jasmine) versus SVs reported in the 1000 Genomes Project (1KGP) after HWE filtering. **b.)** Effect sizes of significant SV-eQTLs mapped from Jasmine SVs or 1KGP SVs. **c.)** CAVIAR posterior probabilities for each gene with significant SV and SNP data. The x-axis is the maximum CAVIAR posterior of a SNP reported as a SNP-eQTL by the GTEx consortium, and the y-axis is the maximum CAVIAR posterior of a Jasmine SV from our mapped SV-eQTLs. Variants above the diagonal line have a higher SV posterior than GTEx SNP posterior. The inset box contains genes with highly causal (posterior >0.8) SVs. **d.)** Jasmine SV distance to the nearest ENCODE cCRE versus CAVIAR posterior. The histogram shows the distribution of distances to ENCODE cCREs. **e.)** Genotype and gene expression distribution in 1KGP samples for novel SEMA5A deletion. **f.)** Manhattan plot for SNPs and the novel SV near SEMA5A, with p value measured by Wilcoxon rank-sum test. The green point is the SNP reported in GTEx eQTLs (chr5_9431336_A_T); other points are colored by LD to that SNP.

In order to evaluate which SVs are likely to have causal effects on their associated genes, we used the fine-mapping tool CAVIAR (Hormozdiari et al. 2014) to measure the posterior probability that any given SV is causal compared to nearby SNPs within a 1 Mbp window, taking into account possible linkage disequilibrium (LD) between variants. We found that SVs had high posterior scores (>0.1) at 68 genes out of 1,863 genes examined (3.65%). Additionally, when compared to existing databases of SNP-eQTLs from the GTEx project (Chiang et al. 2017), SVs had a higher CAVIAR posterior than reported SNPs for 53.5% of genes (**Figure 6c**). This shows that previously undetected SVs are likely causal at a large number of sites where the effects on gene expression were reported as SNP-eQTLs instead.

When examining the CAVIAR posteriors for our data, we found that SVs with higher CAVIAR posteriors are enriched for positions overlapping with or very close to ENCODE candidate cis-regulatory elements (**Figure 6d**), indicating that a number of the high-scoring variants are functionally relevant. We also found that higher CAVIAR posteriors are associated with other regulatory elements, distance to the associated gene (as previously reported in (Chiang et al. 2017)), as well as to FunSeq high occupancy of transcription factor (HOT) regions (Fu et al. 2014) (**Supplementary Figures 24-25**).

Inspecting all SV-gene pairs with a CAVIAR posterior greater than that of any previously reported SNP-eQTL for that gene (and greater than 0.2 overall), we identified several potentially functional SVs in high linkage disequilibrium (LD) with reported SNPs. Among these newly discovered SV-sQTLs is a noncoding deletion associated with the expression of SEMA5A, a gene involved in neural development that has been implicated as an autism susceptibility gene (Melin et al. 2006; Duan et al. 2014) (**Figure 6e**). We found that while a number of SNPs are associated with this gene’s expression, including SNPs reported in the GTEx SNP-eQTL dataset, the most highly associated variant is the structural variant (**Figure 6f**). Other small deletions in SEMA5A have been previously associated with neurodevelopmental disorders (Mosca-Boidron et al. 2016), but this deletion was not previously reported as it is difficult to detect from short-read data alone. This suggests that previous studies exclusively examining SNPs may have ascribed functional relevance to SNP-eQTLs in close LD with the SV. In addition to SEMA5A, we also found several additional examples in LRGUK, CSF2RB, CAMKMT, and several other genes where reportedly functional SNPs are in close LD with potentially more functionally significant SVs, which are underrepresented or ignored in existing eQTL studies (**Supplementary Figures 28-30**).

## Discussion

Here we introduced Jasmine, a fast and accurate method for population-level structural variant comparison and analysis. It improves upon existing methods and achieves highly accurate results by merging pairs of variants in increasing order of their breakpoint distance, while maintaining favorable scaling qualities through the use of a KD-tree to efficiently locate nearby variant pairs. Jasmine also separately processes the SV calls by chromosome and SV type and strand to enable built-in parallelization, while many alternative methods incorrectly combine SVs of different types. By combining Jasmine with additional novel methods and carefully optimizing existing methods, we produced an SV-calling pipeline that reduces the rate of Mendelian discordance by more than a factor of five over prior pipelines, while at the same time being applicable to large cross-technology cohorts and resolving a number of issues encountered when using other methods. Finally, by calling SVs in 31 publicly available long-read samples with our pipeline we developed and released a database of human structural variants. By genotyping these variants in 444 short-read samples from the 1000 Genome Project, we catalogued novel eQTLs across the human genome, including in medically relevant genes.

While Jasmine offers highly accurate population SV analysis, we remain limited by the sequencing data that is available. A major challenge we faced when applying our methods to a cohort consisting of samples from multiple sequencing technologies was the additional noise in the samples sequenced with high-error CLR reads (**Supplementary Figure 18**). While we mitigated this noise through computational means such as double thresholding and carefully tuned parameters, we expect that even more accurate SV calls could be obtained by using HiFi or ONT sequencing for all samples. In addition, there were systematic anomalies in the SV calls in highly repetitive regions such as the centromere and satellite repeats and an overall excess of variants that are found in all samples. There has recently been work to improve the reference genome to more accurately reflect these regions (Nurk et al. 2021), and as tools for aligning to and calling variants in these regions continue to mature, we expect the accuracy of these calls to even further improve. Finally, while we have detected a large number of SVs in the 31 samples we studied, there is still much to be discovered. As the costs of long-read genome sequencing continue to decrease, we expect to apply these methods to even larger populations, as well to other species, to deepen our understanding of the role of SVs in disease, development, and evolution.

## Acknowledgements

We thank Fritz Sedlazeck and Michael Alonge for helpful discussions. This work was supported, in part, by National Science Foundation grants DBI-1350041, IOS-1445025, IOS-1732253 and IOS-1758800 and National Institutes of Health grants NHGRI U24-HG010263 and NIGMS R35GM139580. This work was also supported in part by the Mark Foundation for Cancer Research (19-033-ASP) and a Microsoft Research Fellows award. Part of this research project was conducted using computational resources at the Maryland Advanced Research Computing Center (MARCC).

## Online Methods

### Refined Variant Breakpoints and Sequences with Iris

Many existing long-read SV callers identify variants from read alignments based on signatures such as an extended gap in the alignment or a segment of the read which aligns to a distant region of the genome (Sedlazeck, Rescheneder, et al. 2018; Jiang et al. 2020). In the widely used variant caller sniffles (Sedlazeck, Rescheneder, et al. 2018), a variant is called when multiple reads show similar signatures that cluster together based on their type, span, and location. However, when reporting the variant’s breakpoints and sequence, the alignment from a single representative read (chosen arbitrarily) is used to infer this information. This is particularly problematic for insertions, where the novel sequence being inserted is taken directly from the single read. Since some read technologies, such as CLR and ONT have error rates of 5% or higher, it is expected that the sequence reported will have a sequence with a similar or higher rate of divergence from the true insertion sequence (**Supplementary Figure 1**). When comparing across samples, especially those sequenced with different technologies with different error models, this may cause the same variant in both individuals to be falsely identified as two separate variants.

Addressing this, we introduce Iris, a method for refining the breakpoints and novel sequence of SV calls by aggregating information from multiple reads which support each variant call (**Figure 1**). Iris refines each variant call separately, but supports the processing of multiple variants in parallel. In the case of an insertion variant call, Iris starts with an initial sequence consisting of the variant sequence plus flanking sequence from the reference genome (default 1kb on each side of the variant). Then, it gathers all of the reads which support the variant’s presence - indicated by the RNAMES field in the output of sniffles - and aligns those reads to the initial sequence with minimap2 (Li 2018). These alignments are used as input to the polishing software racon (Vaser et al. 2017), which polishes the initial sequence. Finally, the polished sequence is aligned to the reference with minimap2 and the CIGAR string is parsed to extract the insertion in the polished sequence relative to the reference which most closely resembles the original insertion call. If such an insertion is found, the variant call is refined to reflect the updated sequence and breakpoints. Iris also supports the refinement of deletion breakpoints, which is of particular interest when the sequencing technology being used has a biased error model in favor of either insertions and deletions. These are handled similarly to insertions, with the initial sequence instead consisting of the concatenation of the reference sequences immediately before and after the deleted region. Iris is available as a standalone tool at https://github.com/mkirsche/Iris.

#### Simulation Results

To test the performance of Iris on simulated data, we simulated 400 indels with uniformly random lengths - 200 with length [50, 200] and 200 with length [900, 1100] - in a 5 Mbp segment of chr1 (chr1:20000000-24999999). Then, we used SURVIVOR (Jeffares et al. 2017) with a read error and length model trained on HG002 Oxford Nanopore reads to simulate 30x coverage of long reads. We aligned these reads back to the unmodified segment of chromosome 1 with winnowmap (C. Jain et al. 2020) and called SVs with sniffles (Sedlazeck, Rescheneder, et al. 2018). From the insertion SV calls, we measured the similarity scores of the reported sequences to the ground truth using the formula: Similarity(*S, T*) = (1 - EditDistance(*S, T*) / max(length(*S*), length(*T*)). We also refined these variant calls with Iris and measured the similarity score of the updated insertion sequences (**Supplementary Figure 2a**). The average sequence similarity score increased from 94.7% to 98.6%, demonstrating that Iris refinement significantly improves insertion sequence accuracy.

#### Real Results in HG002

While this simulated experiment demonstrated that Iris is able to improve sequence accuracy in simulation conditions, we wanted to ensure that it also improves the novel sequences of true genomic variants, where the novel sequences are typically more repetitive and the alignments noisier than when the insertions are random basepairs. To do this, we used the cell line HG002, which was sequenced with multiple technologies, notably including both ONT and HiFi. While the ONT reads have a high error rate around 8%, the HiFi reads have approximately 99.5% accuracy (Wenger et al. 2019), so even novel insertion sequences taken from only a single HiFi read are expected to be highly accurate. Therefore, we used winnowmap and sniffles for variant calling as in the simulated experiment, but used the HiFi SV calls’ sequences in place of a ground truth. For each ONT SV call, we matched it with the nearest HiFi call if it was within 1 kbp, they shared at least 50% sequence identity, and no other ONT call had already matched with it. This resulted in 13,467 matched ONT calls before and 14,401 after refinement, with 12,978 having a matching HiFi call both before and after refinement. Among these, 9,522 (73.37%) had been changed by Iris. The average sequence identity among these 9,522 SVs increased from 91.6% before Iris to 96.2% after Iris, and the distributions of sequence accuracy scores are shown in **Supplementary Figure 2b**.

### Comparing Variant Calls at Population Scale with Jasmine

In order to perform SV inference at population scale and identify variants associated with diseases or phenotypes, it is important to identify when multiple individuals share the same (or functionally identical) variants. However, the same variant call can manifest differently in unique samples because of sequencing error or samples being processed with different sequencing technologies, levels of coverage, or upstream alignment and variant calling software. These differences, along with the increasing availability of long-read sequencing data for many individuals, highlight the need for careful variant comparison when analyzing SVs in multiple samples.

We refer to the problem of consolidating multiple variant callsets into a single set of variants as the “SV merging problem”. This is because the problem consists of identifying variant calls in separate samples which correspond to the same variant and merging them into a single call which is annotated with the samples in which it is present. A number of methods already exist for SV merging, but each has major issues such as invalid merges, results which vary significantly based on the order of input samples, or high levels of Mendelian discordance when evaluated on trio datasets.

#### Jasmine Methods

We introduce Jasmine, a novel method which solves the SV merging problem. Jasmine takes as input a list of VCF files consisting of the variant callsets for each individual, and produces a single VCF file in which each variant is annotated with a list of samples in which it is present (as well as the IDs of the input calls which correspond to that variant).

Jasmine first separates the variants by their chromosome (or chromosome pair in the case of translocations), variant type, and strand. Each of these groups is processed independently with an option for parallelization because no two variants in different groups could be representations of the same variant. When processing a group of variants, Jasmine represents each variant as a 2-D point in space representing the start position and length of the variant. When represented this way, variants which are closer together along the genome (and are therefore more likely to represent the same variant) have a smaller Euclidean distance between them. Consequently, each pair of variants can be assigned a quantitative distance which reflects how dissimilar they are.

After projecting these variants into 2-D Euclidean space, Jasmine implicitly builds a variant proximity graph, or a graph in which nodes are individual variants and each pair of variants has an edge between them with a weight corresponding to the Euclidean distance between them. Then, the SV merging can be framed as selecting a set of edges (merges) making up a forest which is a subgraph of the variant proximity graph, and which minimizes the sum of edge weights chosen subject to a few constraints:

1. ***No intra-sample merging:*** No connected component of the forest contains multiple variants from the same individual because they have already been identified as different variants
2. ***Distance threshold:*** No chosen edge has a weight greater than the user-chosen distance threshold (default max(100bp, 50% of variant length))
3. ***Maximality:*** To prevent the trivial solution of no edges, we require that given a set of chosen edges, no additional edges can be added to the solution while still satisfying the other constraints.

Jasmine seeks to solve this optimization problem with a greedy algorithm similar in design to Kruskal’s algorithm for finding a minimum spanning tree. In this algorithm, the set of chosen edges is initially empty, and each edge is considered in order of non-decreasing edge weight. If adding the edge to the solution would violate any of the above constraints given the previously added edges, it is ignored; otherwise, it is added to the solution. When the edges being considered start to exceed the distance threshold, the algorithm terminates.

One issue with this algorithm is that in order to sort the edges by weight, they may need to be loaded into memory. This is problematic because some population datasets, with tens to hundreds of thousands of SVs per sample, include millions of variants, with the number of edges potentially scaling quadratically with the variant count. This is prohibitive even with existing datasets, and will only be more of a problem as even larger datasets are produced. Therefore, Jasmine instead stores the edges implicitly, making use of a KD-tree to quickly find the next smallest edge in the variant proximity graph.

To avoid storing the entire graph in memory, Jasmine maintains a list of a small number of nearest neighbors (initially 4) for each node, which are computed by forming a KD tree with all of the variant points, a data structure which supports k-nearest neighbor queries with a logarithmic runtime with respect to the number of variants. Then, the edge to the single nearest neighbor of each variant is stored in a minimum heap, and it is guaranteed that the first entry removed from this heap will be the edge with the smallest weight in the entire graph. When an edge is processed, the node for which it was the minimum-weight incident edge has its next nearest neighbor added to the heap based on the next entry in its nearest neighbor list. If the list of nearest neighbors for a node becomes empty, the KD-tree is queried for a set of twice as many nearest neighbors, and the list is refilled. In this manner, the next smallest edge in the graph will always be the edge removed from the heap, and the data structures Jasmine uses help to maintain this property without requiring a prohibitively large amount of time or memory. The pseudocode for this algorithm can be found in **Supplementary Note 1**.

#### Jasmine Distance Threshold

When merging variants, it is important to determine for a given variant pair whether or not the two variants are sufficiently close together in terms of their breakpoints to be considered the same variant. In Jasmine, this is based on a distance threshold - if the distance between them (according to the chosen distance metric) is above the threshold they will be considered two different variants, while if their distance is less than or equal to the threshold they will be a candidate for merging. Jasmine offers a number of classes of distance thresholds, including constant thresholds, thresholds which vary based on a fixed proportion of each variant’s size, or even per-variant distance thresholds. By default, the distance threshold for Jasmine is max(100bp, 50% of variant length). We measured the difference in merging when using different values for the *min_dist* parameter, which is 100 by default, (**Supplementary Figure 3)**, and found that while larger values for this parameter offer lower Mendelian discordance, these more lenient thresholds perform poorly in a cross-technology cohort setting because of false merges, and 100bp offers a good balance in performance across use cases.

### Building an SV Inference Pipeline

Our SV inference pipeline is implemented in Snakemake, and supports multithreaded as well as multi-node execution. It takes as input a list of FASTQ files for each sample being studied as well as a reference genome, and produces as its final output a VCF file containing population-level SV calls. It is highly customizable, supporting unique configurations for alignment and variant calling on a per-sample or per-sequencing-technology level. It also enables the user to specify the alignment software to use - ngmlr, winnowmap, and minimap2 - and separately sets recommended default parameters for samples sequenced with each specific technology. On each sample we processed, the pipeline took about a day to run on a single Intel Cascade Lake 6248R compute node with 48 cores and 192GB RAM at 3.0GHz. The Snakemake files to run the pipeline are included in the Jasmine repository: https://github.com/mkirsche/Jasmine/tree/master/pipeline.

### Evaluating Mendelian Discordance

When performing *de novo* variant analysis, we are particularly interested in Mendelian discordant variants, or variants which are called as present in the child of a trio but neither parent. This includes genuine *de novo* variants, but in practice most of these calls are actually false *de novo* variants caused by errors in variant calling or merging. Accordingly, one major goal of trio SV inference is to reduce the number of discordant variants while retaining any true *de novo* variants in that set.

To measure Mendelian discordance, we called variants in the Ashkenazim individual HG002 as well as their parents HG003 (46,XY) and HG004 (46,XX). We merged these three callsets with Jasmine (or other merging software when comparing them to Jasmine), and counted the number of variants which were identified in HG002 but not merged with any variants from either parent. We then filtered these variants by confidence by requiring that they be supported by at least min(10, 25% of average coverage) of the reads and have a length of at least 30. In addition, we filtered out any variants which were not marked with the PRECISE INFO field by the sniffles variant calling. The discordance rate was calculated as the quotient of the number of discordant variants over the total number of variants in the merged and filtered trio callset.

### Optimized Sniffles Variant Calling Parameters

Similar to the HiFi analysis in **Figure 2c**, we used Mendelian discordance to measure the effects of different variant calling parameters in CLR data for HG002. We varied the *max_dist* parameter when running sniffles for variant calling and measured the number of variants and discordance for each trio callset. **Supplementary Figure 4** shows the effect of this parameter on these metrics, and based on these results we used *max_dist*=50 for calling variants from CLR data.

Next, to optimize variant calling parameters in ONT data from HG002, we repeated the experiment used for CLR data, varying the *max_dist* variant calling parameter in Sniffles and measured the number of variants and discordance for each trio callset. These results are shown in **Supplementary Figure 5**, and based on them we used *max_dist*=50 for calling variants from ONT data. While this doesn’t give the lowest discordance rate, all settings examined yielded less than 1% discordance, so we used a value of 50 to enable a high degree of variant discovery and consistency across technologies.

### Double Thresholding

To reduce the impact of threshold effects on variant calling, our pipeline uses two different variant calling thresholds: a highly specific, strict high-confidence threshold and a highly sensitive, more lenient low-confidence threshold. To be a high-confident call, a variant must be at least 30bp long supported by a number of reads greater than or equal min(10, 25% of average coverage over that sample); otherwise a variant is called with low confidence if it is at least 20bp long and supported by at least two reads. All of the variants that meet either threshold are used as input to Jasmine’s cross-sample merging, and any low-confidence variants that do not get merged with any high-confidence variants are discarded. This allows variants which are close to the strict threshold to be properly detected in all of the samples in which they are present (**Supplementary Figure 6**).

### Associating Structural Variants to Genes

To obtain genotypes for SV-gene association, we called SVs in 31 long-read samples with our inference pipeline and merged them into a unified cohort-level callset with Jasmine. We then genotyped these SVs with Paragraph after filtering out translocations and other variants which Paragraph cannot genotype, for a total of 189,581 genotyped variants across 444 individuals. Following previous studies (Chen et al. 2019), we then used the Hardy-Weinberg Equilibrium (HWE) test to filter out variants not consistent with population genetic expectations, removing variants found to be significant with p < 0.0001 using an exact test of HWE (Wigginton, Cutler, and Abecasis 2005). After filtering with HWE and additionally removing any variants that were left uncalled in 50% or more of the samples, we were left with 138,715 variants across the 444 individuals (**Supplementary Figure 23**).

We examined common *cis*-SV-eQTLs by associating our SV genotypes to gene expression data in the same cell lines collected by the GEUVADIS consortium (Lappalainen et al. 2013). We first paired each gene with every structural variant that has a MAF ≥0.05 and resides within a window of 1 Mbp from the gene’s TSS. We then tested whether the distribution of normalized (zero-mean, unit variance) gene expression is different for those individuals with or without the variant by using a Wilcoxon rank-sum test for each variant-gene pair with a p-value cutoff reflecting a Benjamini-Hochberg multiple testing correction with an FDR of 0.1. After identifying a set of significantly-associated SV-eQTLs, we fit a linear model between each variant genotype (where reference is encoded as 0 and the alternate allele is encoded as 1 if heterozygous and 2 if homozygous) and gene expression in order to determine the effect size (β) and the R^2^ of the association. We then analyzed the relationship between the effect size and various features of the SV or gene.

#### Comparing SVs and SNP-eQTLs with Fine Mapping

We used the dataset of SNP-eQTLs from the GTEx project for all tissues (Chiang et al. 2017) as a set of known SNP-eQTLs which we could use as a benchmark to compare the effects of SVs to SNPs on genes for which both may be associated. We examined the set of genes for which there were both associated SNP-eQTLs in GTEx (which were also significantly associated in our data) and significantly-associated SVs from our callset within a 1MB window. We then collected a set of 1000 most-closely associated variants (SNP or SV) to each gene within the 1MB window and computed the Z-score from a linear regression as well as the linkage disequilibrium between each pair of variants. We used these values as input to the fine-mapping program CAVIAR (Hormozdiari et al. 2014) in order to predict which variants within the set are causal. We used CAVIAR’s posterior probability as a measure of how likely a particular variant was to be causal.

#### Measuring Enrichment of SVs based on CAVIAR Scores

We examined the relationship between CAVIAR’s posterior probability for each SV’s most highly associated gene and various variant features, such as the distance to various regulatory elements (**Supplementary Figure 25**). We used the `bedtools closest` function to compute the distance between each SV and the nearest ENCODE candidate cis-regulatory element from the UCSC genome browser (Navarro Gonzalez et al. 2021) (**Supplementary Figure 25a**). Using the Ensembl Regulatory Build (Zerbino et al. 2015), we performed a similar distance calculation to measure the distance between each variant and the nearest Ensembl Regulatory Element (**Supplementary Figure 25b**).

We also examined the relationship between CAVIAR posterior probability and various conservation scores, as well as other sequence features such as GC content. To compute conservation scores, inspired by previous works (Abel et al. 2020), we used pyBigWig to extract regions covered by the SV and computed the mean of the top 10 scores of individual bases within that region. For insertion variants, we extracted the flanking reference sequence - 75 basepairs in each direction - to assess the conservedness of the affected context. We calculated CADD scores (Rentzsch et al. 2019), LINSIGHT scores (Huang, Gulko, and Siepel 2017), and PhastCons (Hubisz, Pollard, and Siepel 2011) in a similar fashion. Based on these prediction scores, we do not observe signs of enrichment of extreme pathogenicity or conservation among SVs with high CAVIAR posteriors (**Supplementary Figures 26-27**). We also do not observe a pattern among the GC percentage for SVs with high CAVIAR posteriors (**Supplementary Figure 27a**). However, larger-scale studies are needed to make definitive conclusions, as the number of SVs we observed with high CAVIAR posterior are limited.

## Data Availability

The sequencing data used in this study is available from the publications listed in Supplemental Table 1 and Supplemental Table 2. All variant calls are available at http://data.schatz-lab.org/jasmine/.

