## Supplemental Materials for "Jasmine: Population-scale structural variant comparison and analysis"

**Supplemental Table 1. Data Used for Trio Analysis.** 2
**Supplemental Table 2. Data Used for Cohort Analysis.** 3
**Supplemental Table 3. Software Versions.** 4
**Supplementary Figure 1. Variable Breakpoints and Sequences among Individual Reads.** 5
**Supplementary Figure 2. Improved Insertion Sequence Accuracy with Iris.** 6
**Supplementary Figure 3. Mendelian Discordance across Jasmine Distance Thresholds in the HG002 Trio.** 7
**Supplementary Figure 4. Optimized Variant Calling Parameters for CLR.** 8
**Supplementary Figure 5. Optimized Variant Calling Parameters for ONT.** 9
**Supplementary Figure 6. Double Thresholding to Reduce Threshold Effects.** 10
**Supplementary Figure 7. Discordance by Read Support with Double Thresholding.** 11
**Supplementary Figure 8. Length and Read Support among HG002 HiFi Variants with Double Thresholding.** 12
**Supplementary Figure 9. PCA of 1KGP SV Genotypes.** 13
**Supplementary Figure 10. Read Support across Sequencing Technologies.** 14
**Supplementary Figure 11. Distribution of SV Type and Length in HG002 HiFi Trio by Genomic Context.** 15
**Supplementary Figure 12. HG002 HiFi Trio Sample Presence by Genomic Context.** 16
**Supplementary Figure 13. HG002 HiFi Trio Cross-Technology Agreement by Genomic Context.** 17
**Supplementary Figure 14. Additional Potential de novo SVs in HG002.** 18
**Supplementary Figure 15. Additional Potential de novo SVs in HG002.** 19
**Supplementary Figure 16. Lower-Confidence Potential de novo SV in HG002.** 20
**Supplementary Figure 17. Variant Counts per Sample.** 21
**Supplementary Figure 18. Variant Counts per Sample including Low-Confidence SVs.** 22
**Supplementary Figure 19. Allele Frequencies of all Merging Software.** 23
**Supplementary Figure 20. Length-Filtered Allele Frequencies of all Merging Software.** 24
**Supplementary Figure 21. Example of Breakpoint Directionality.** 25
**Supplementary Figure 22. HiFi Alignments for Detecting SEMA5A-Associated SV.** 26
**Supplementary Figure 23. 1KGP Allele Frequencies before and after HWE Filtering.** 27
**Supplementary Figure 24. SV Caviar Posteriors by HOT Region and Gene Distance.** 28
**Supplementary Figure 25. SV CAVIAR Score vs. Distance to Regulatory Elements.** 29
**Supplementary Figure 26. SV CAVIAR Score vs. CADD and PhastCons Scores.** 30
**Supplementary Figure 27. SV CAVIAR Score vs. GC Content and LINSIGHT Score.** 31
**Supplementary Figure 28. Potential Functionally Relevant Deletion in LRGUK.** 32
**Supplementary Figure 29. Potential Functionally Relevant Insertion in CSF2RB.** 33
**Supplementary Figure 30. Potential Functionally Relevant Insertion in CAMKMT.** 34
**Supplemental Note 1. Jasmine Merging Algorithm.** 35
**Supplemental Note 2. Commands Used.** 37

**Supplemental Table 1. Data Used for Trio Analysis.**

| Sample | HiFi Coverage | ONT Coverage | CLR Coverage |
|---|---|---|---|
| HG002 | 35.2499 | 46.6151 | 54.8693 |
| HG003 | 33.6795 | 80.6551 | 25.6278 |
| HG004 | 33.1812 | 83.1599 | 23.4694 |

Source: https://ftp-trace.ncbi.nlm.nih.gov/ReferenceSamples/giab/data/AshkenazimTrio/

**Supplemental Table 2. Data Used for Cohort Analysis.**

| Tech | Sample | Coverage | Study | Ancestry |
|---|---|---|---|---|
| HiFi | HG001 | 29.4987 | GIAB | CEU |
| HiFi | HG00512 | 29.3707 | 1KGP | CHS |
| HiFi | HG00513 | 40.3823 | 1KGP | CHS |
| HiFi | HG006 | 32.4010 | GIAB | CHS |
| HiFi | HG00731 | 32.9366 | 1KGP | PUR |
| HiFi | HG00732 | 21.2571 | 1KGP | PUR |
| HiFi | HG007 | 36.1509 | GIAB | CHS |
| HiFi | HG01109 | 31.7902 | HPRC | PUR |
| HiFi | HG01243 | 34.8145 | HPRC | PUR |
| HiFi | HG01442 | 36.9866 | HPRC | CLM |
| HiFi | HG02055 | 39.0903 | HPRC | ACB |
| HiFi | HG02080 | 33.7257 | HPRC | KHV |
| HiFi | HG02109 | 30.2620 | HPRC | ACB |
| HiFi | HG02145 | 35.7587 | HPRC | ACB |
| HiFi | HG02723 | 45.4921 | HPRC | GWD |
| HiFi | HG03098 | 35.1080 | HPRC | MSL |
| HiFi | HG03492 | 33.2615 | HPRC | PJL |
| HiFi | NA19238 | 24.9931 | 1KGP | YRI |
| HiFi | NA19239 | 25.8028 | 1KGP | YRI |
| ONT | HG003 | 80.6551 | GIAB | ASH |
| ONT | HG004 | 83.1599 | GIAB | ASH |
| CLR | AK1 | 79.2865 | Audano | EAS |
| CLR | CHM13 | 97.1029 | Audano | EUR* |
| CLR | CHM1 | 51.2768 | Audano | EUR* |
| CLR | HG00268 | 69.5876 | Audano | FIN |
| CLR | HG01352 | 56.2097 | Audano | CLM |
| CLR | HG02059 | 63.9237 | Audano | KHV |
| CLR | HG02106 | 59.9712 | Audano | PEL |
| CLR | HG04217 | 128.5960 | Audano | ITU |
| CLR | HX1 | 76.6489 | Audano | EAS |
| CLR | NA19434 | 58.3505 | Audano | LWK |

*Hydatidiform mole for which only the super-population has been reported

1KGP: http://ftp.1000genomes.ebi.ac.uk/vol1/ftp/data_collections/HGSVC2/working/
GIAB: http://ftp-trace.ncbi.nlm.nih.gov/giab/ftp/data
HPRC: https://github.com/human-pangenomics/HPP_Year1_Data_Freeze_v1.0
Audano: (Audano et al. 2019)

**Supplemental Table 3. Software Versions.**

| Software | Version |
|---|---|
| Jasmine | 1.1.0 |
| Iris | 1.0.4 |
| sniffles | 1.0.11 |
| winnowmap | 2.0 |
| racon | 1.4.10 |
| minimap2 | 2.17 |
| samtools | 1.9 |
| SURVIVOR | 1.0.7 |
| svtools | 0.5.1 |
| svimmer | 0.1 |
| dbsvmerge | commit 85b3687a54ce21ba25862c58707daa212b9fbcbd |
| svpop | commit 8be50c55f8e81f8c701077bb9c00ee5bea3e0d2b |
| Paragraph | 2.4 |
| CAVIAR | commit 135b58baffac92b5e9b45f8db78315a9b4d713bc |
| plink | 1.90b6.4 |
| snphwe | 1.0.2 |

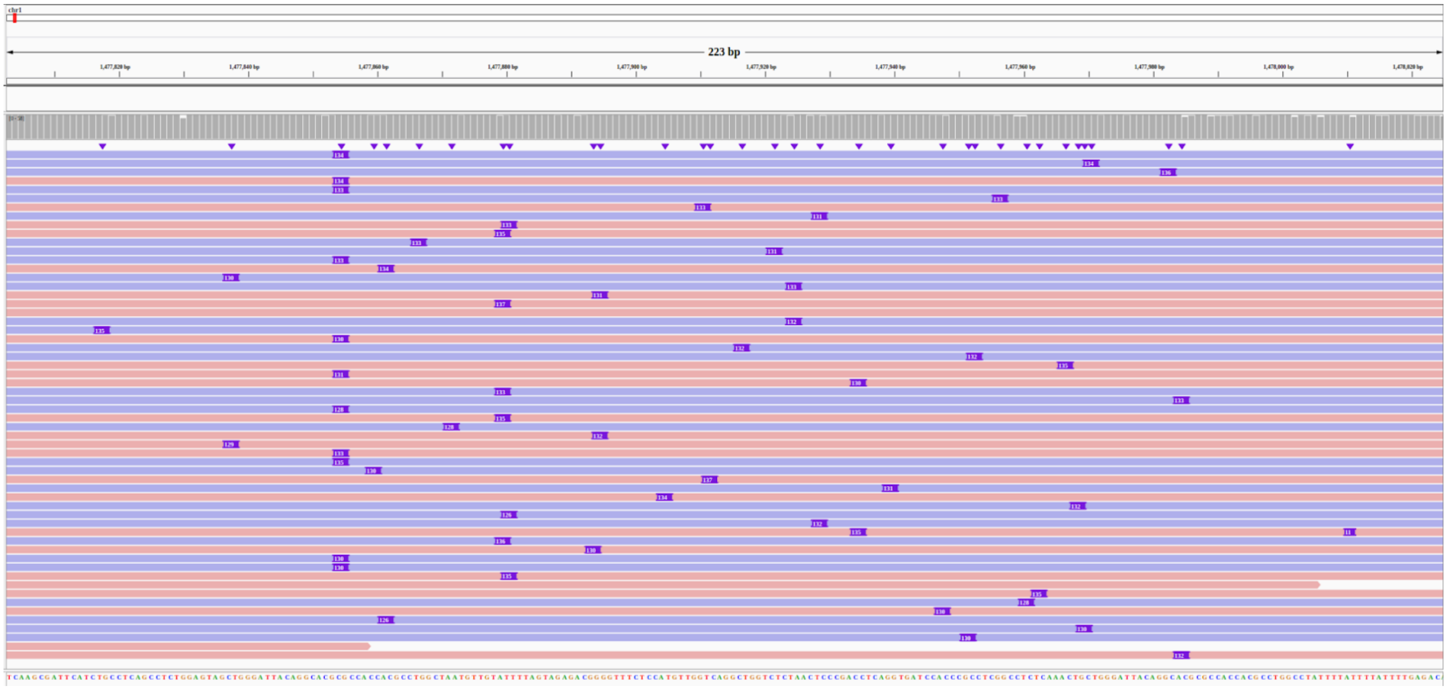


**Supplementary Figure 1. Variable Breakpoints and Sequences among Individual Reads.**
This figure shows an insertion SV in HG002 at chr1:1477881 in which the breakpoints and sequence length vary among individual ONT reads.

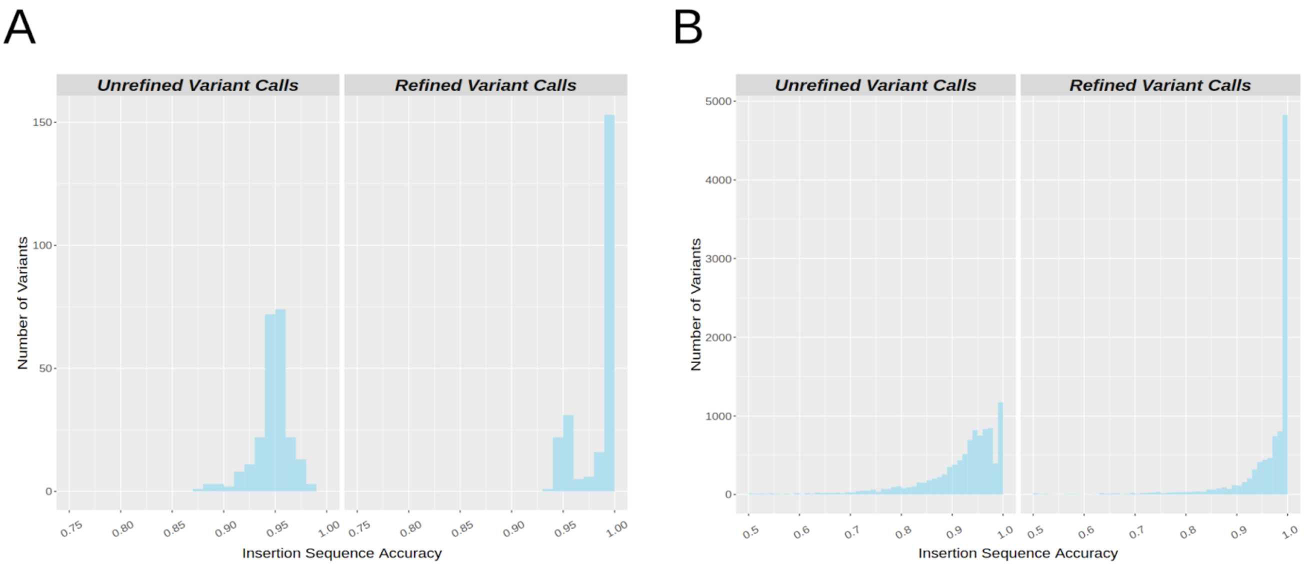


**Supplementary Figure 2. Improved Insertion Sequence Accuracy with Iris.**
**a.)** The distribution of insertion sequence accuracy in 200 SV calls from the simulation of human chromosome 1 with and without Iris refinement.
**b.)** The distribution of insertion sequence accuracy in the HG002 SV calls derived from ONT reads, using the HiFi calls as ground truth, with and without Iris refinement.

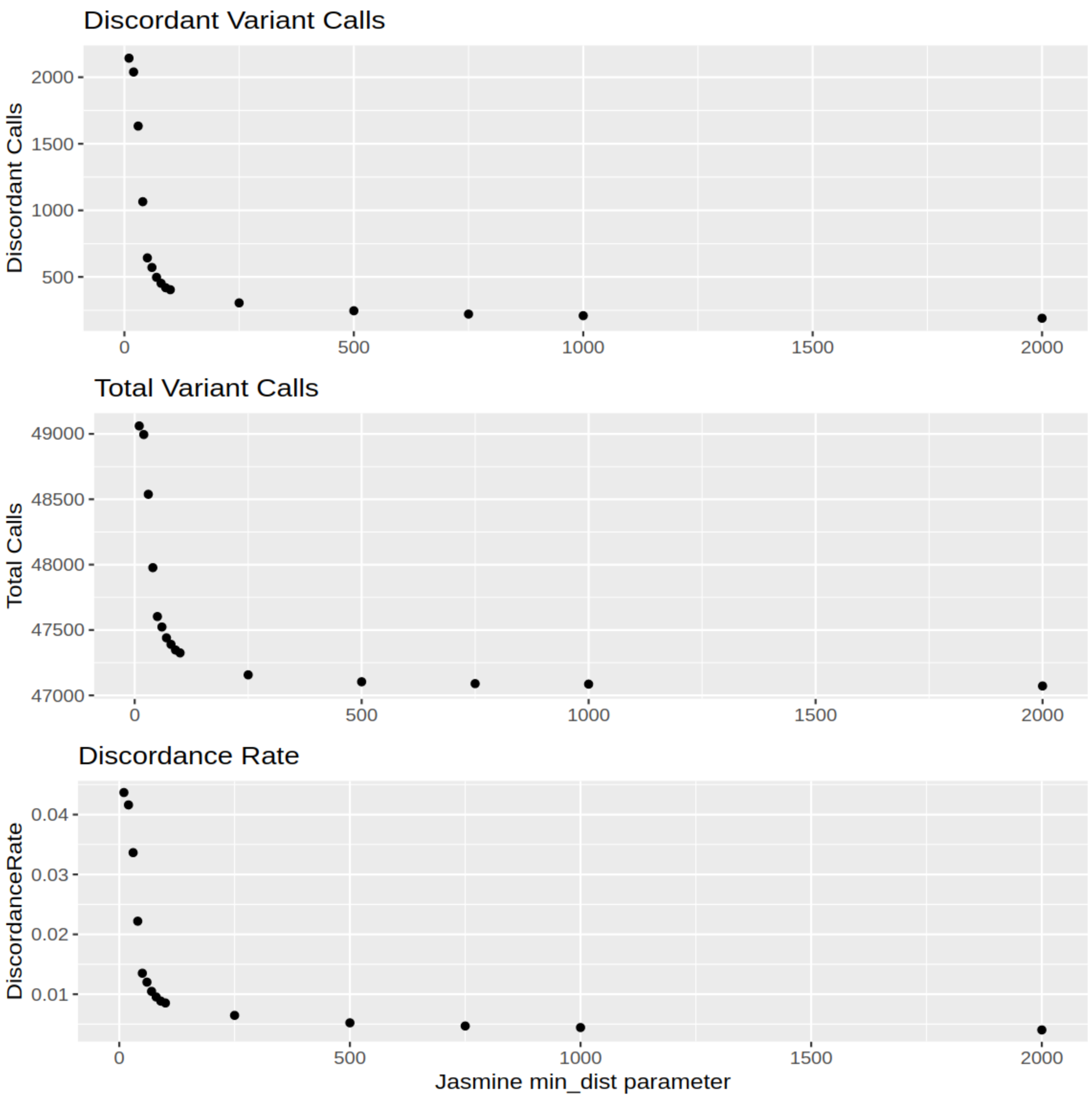


**Supplementary Figure 3. Mendelian Discordance across Jasmine Distance Thresholds in the HG002 Trio.**

We varied the *min_dist* parameter when merging variants in HG002, HG003, and HG004, and observed the total number of variants, the number of discordant variants, and the discordance rate for each run.

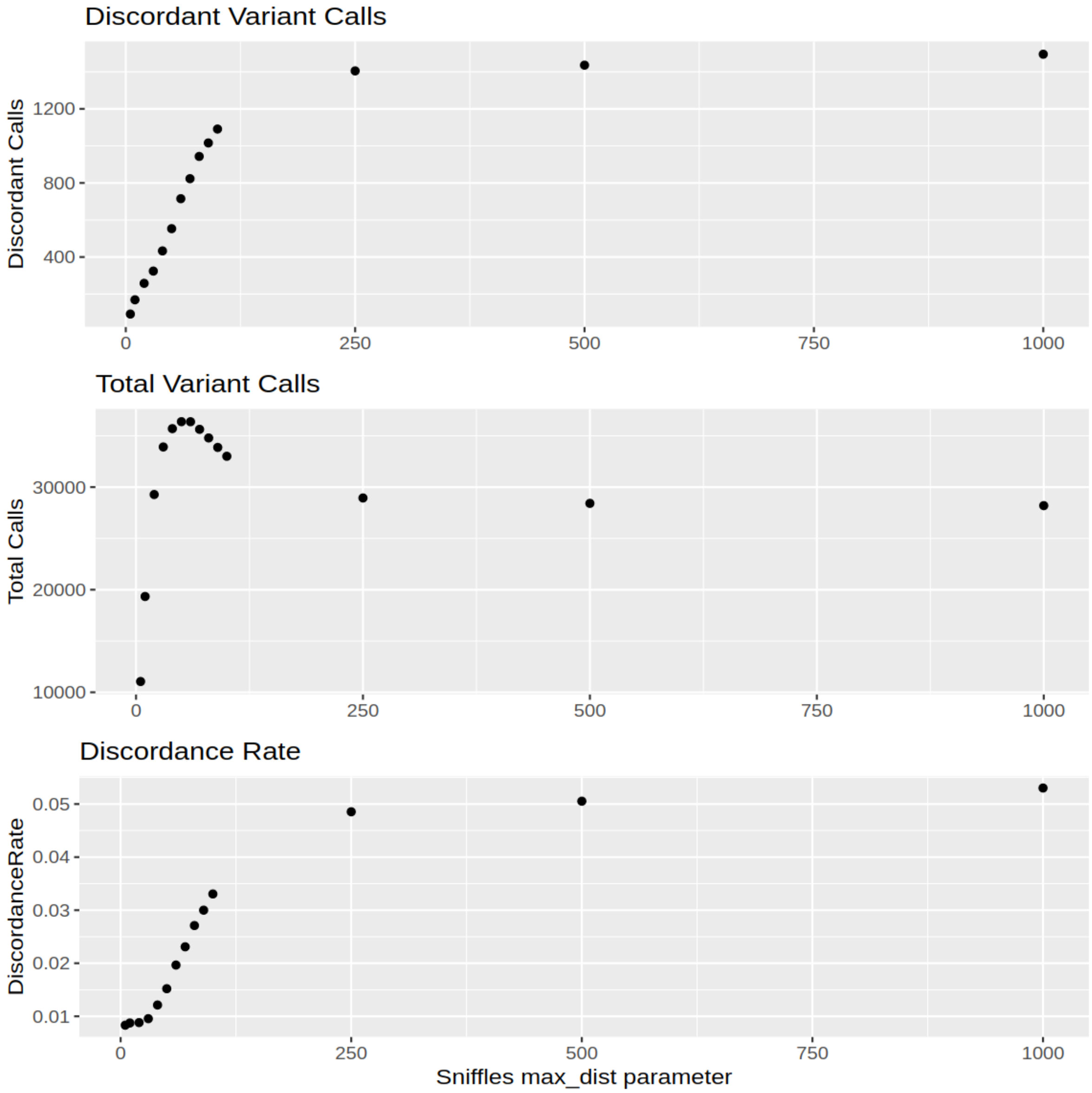


**Supplementary Figure 4. Optimized Variant Calling Parameters for CLR.**
We called SVs in HG002, HG003, and HG004 from CLR reads using different values of the *max_dist* parameter in sniffles and merged each trio callset with Jasmine. For each *max_dist* value we measured the total number of SVs in the trio, the number of discordant variants, and the discordance rate.

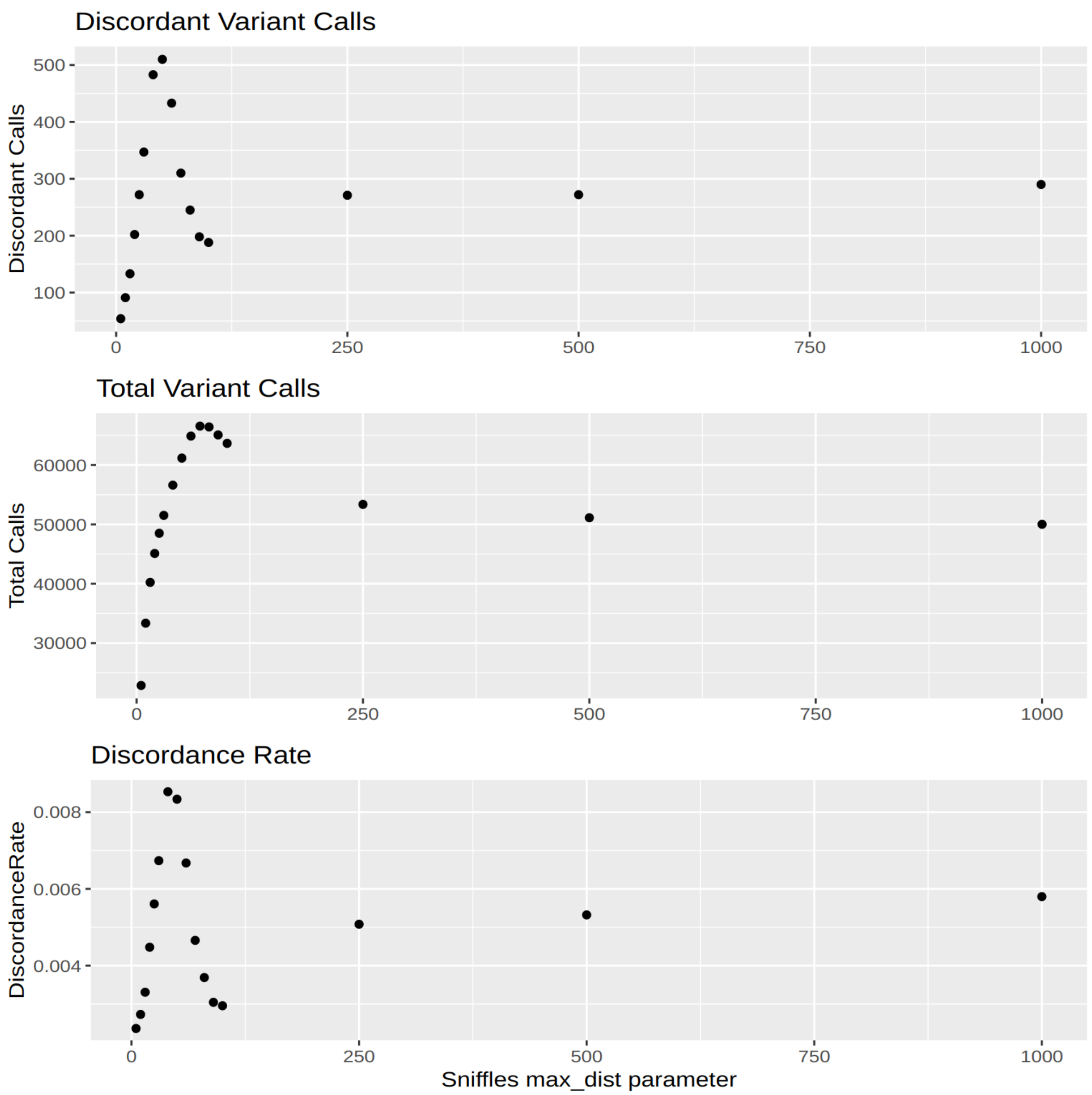


**Supplementary Figure 5. Optimized Variant Calling Parameters for ONT.**
We called SVs in HG002, HG003, and HG004 from ONT reads using different values of the *max_dist* parameter in sniffles and merged each trio callset with Jasmine. For each *max_dist* value we measured the total number of SVs in the trio, the number of discordant variants, and the discordance rate.

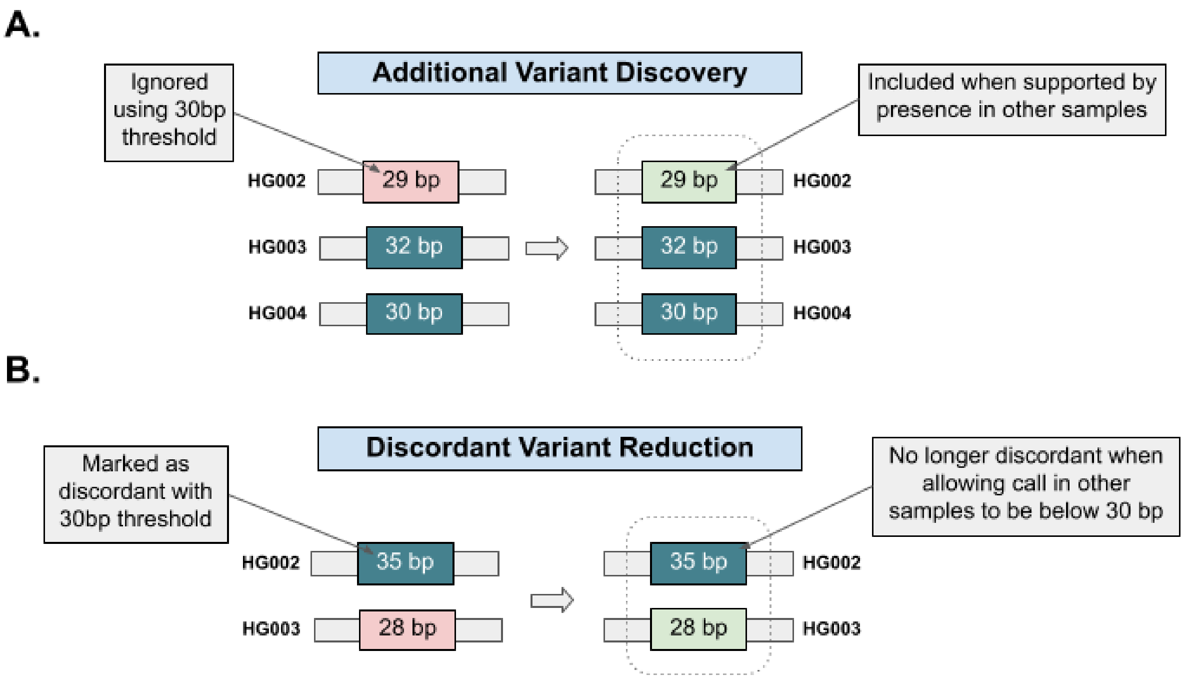


**Supplementary Figure 6. Double Thresholding to Reduce Threshold Effects.**
To avoid cases where variants with length or read support near the variant calling threshold are detected in some but not all samples where they are present, we use a double threshold. In the case of trio analysis, we are able to both **a.)** discover more variants in the child and **b.)** reduce the number of discordant variants compared to using a single threshold.

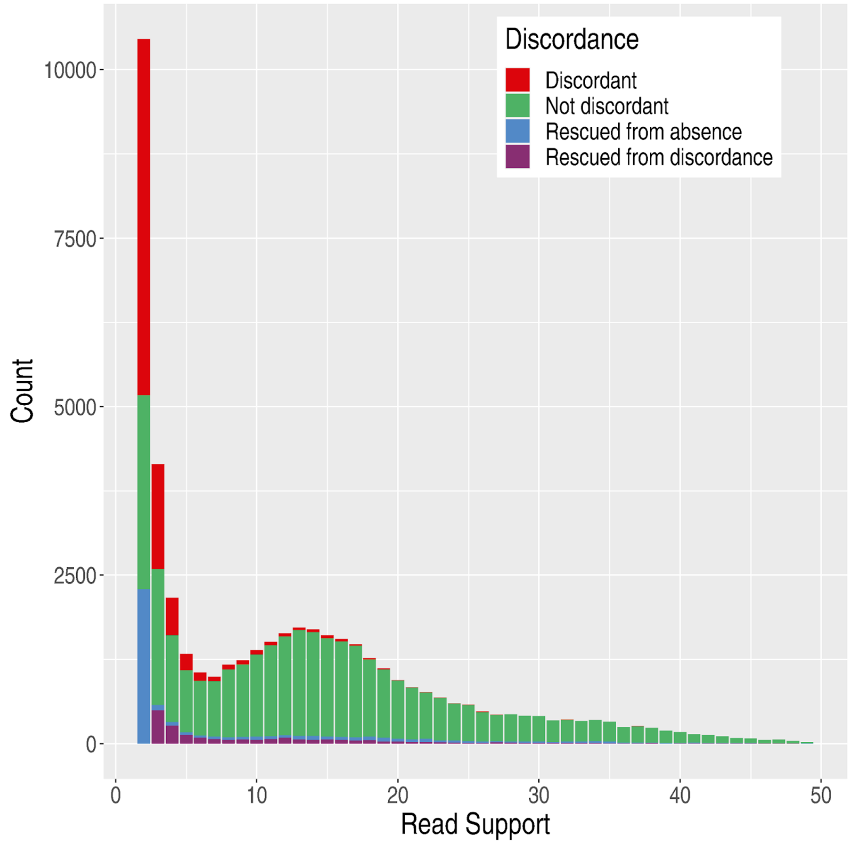


**Supplementary Figure 7. Discordance by Read Support with Double Thresholding.**
This illustrates the read support distribution of SVs in HG002 called from HiFi data. As the read support increases, the number of discordant variants decreases, but even among variants with low read support there are many which are not discordant, and double thresholding improves our ability to resolve them.

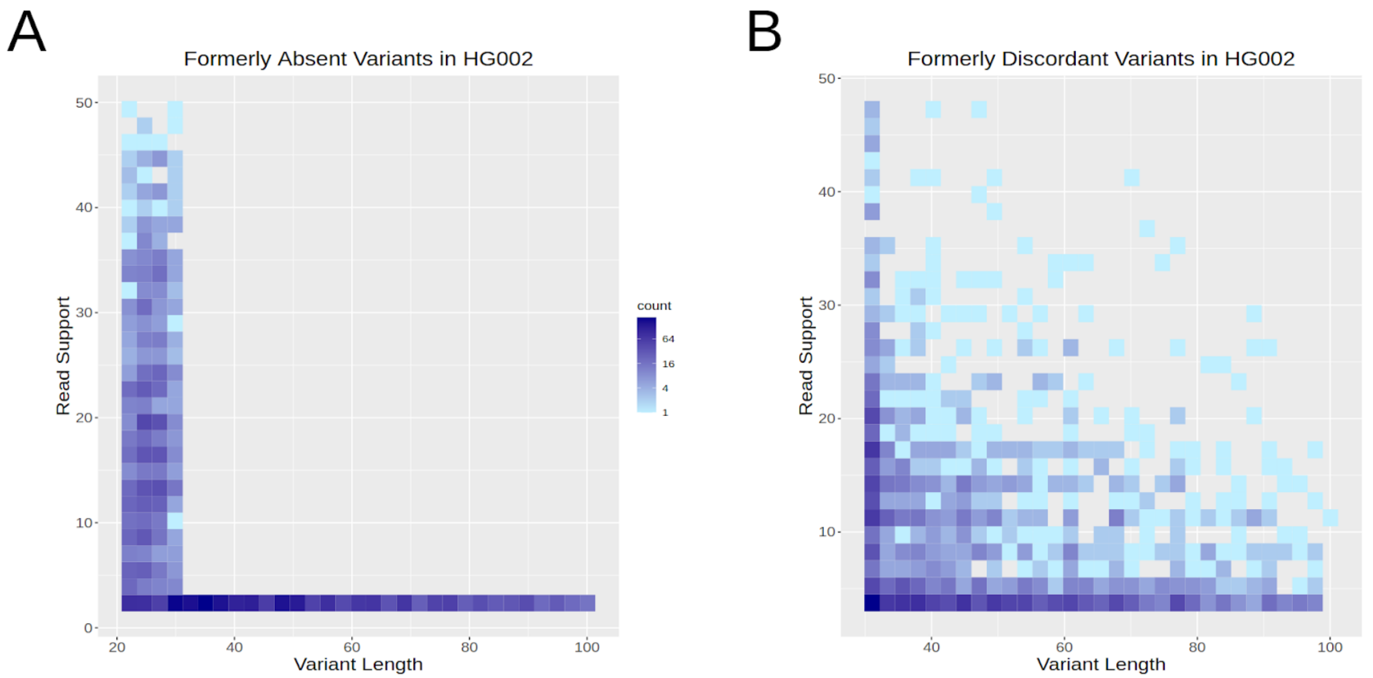


**Supplementary Figure 8. Length and Read Support among HG002 HiFi Variants with Double Thresholding.**
**a.)** The length and read support of variants which would have missed in HG002 if using only a single threshold.
**b.)** The length and read support of variants which would have been discordant in HG002 if using only a single threshold.

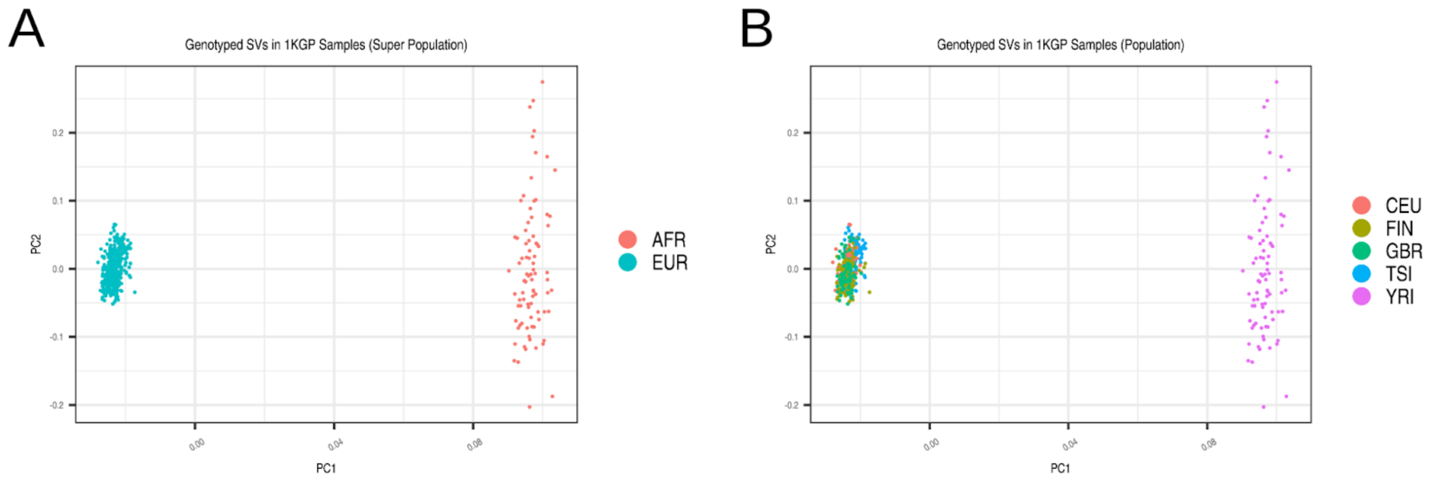


**Supplementary Figure 9. PCA of 1KGP SV Genotypes.**
A two-dimensional projection of the absence/presence vectors of all genotyped SVs in the 444 samples from the 1000 Genomes Project that we used for eQTL analysis, colored by **a.)** superpopulation and **b.)** population.

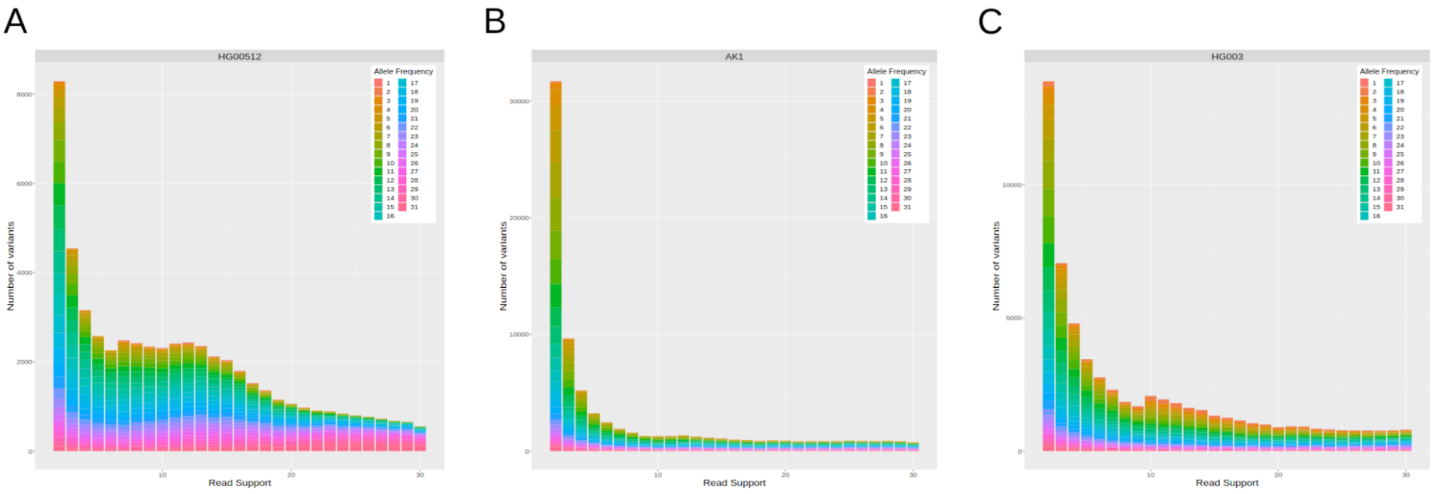


**Supplementary Figure 10. Read Support across Sequencing Technologies.**
In our population analysis, to examine the SV caller's ability to detect reads supporting SVs across technologies, we measured the read support of each SV for a representative sample sequenced with each technology. **a.)** HG00512, a HiFi sample with 29x average coverage. As expected, we see a sharp decrease at about 50% read support, corresponding to the transition from heterozygous to homozygous variants. **b.)** AK1, a CLR sample with 79x average coverage. In this sample, there are many variants supported by only a couple of reads, which is an artifact of reads having high sequencing error and moderate length. **c.)** HG003, an ONT sample with 81x average coverage.

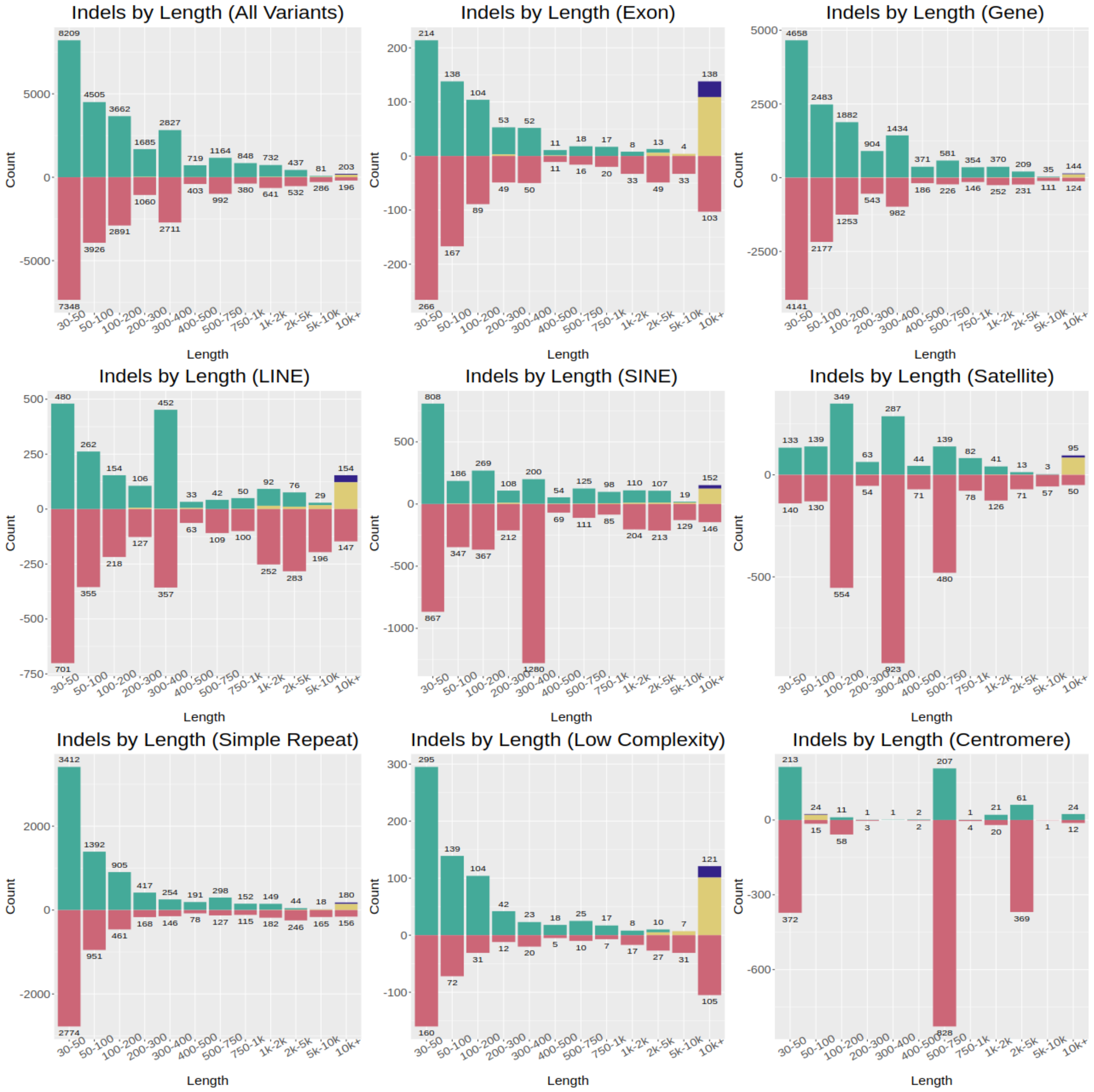


**Supplementary Figure 11. Distribution of SV Type and Length in HG002 HiFi Trio by Genomic Context.**

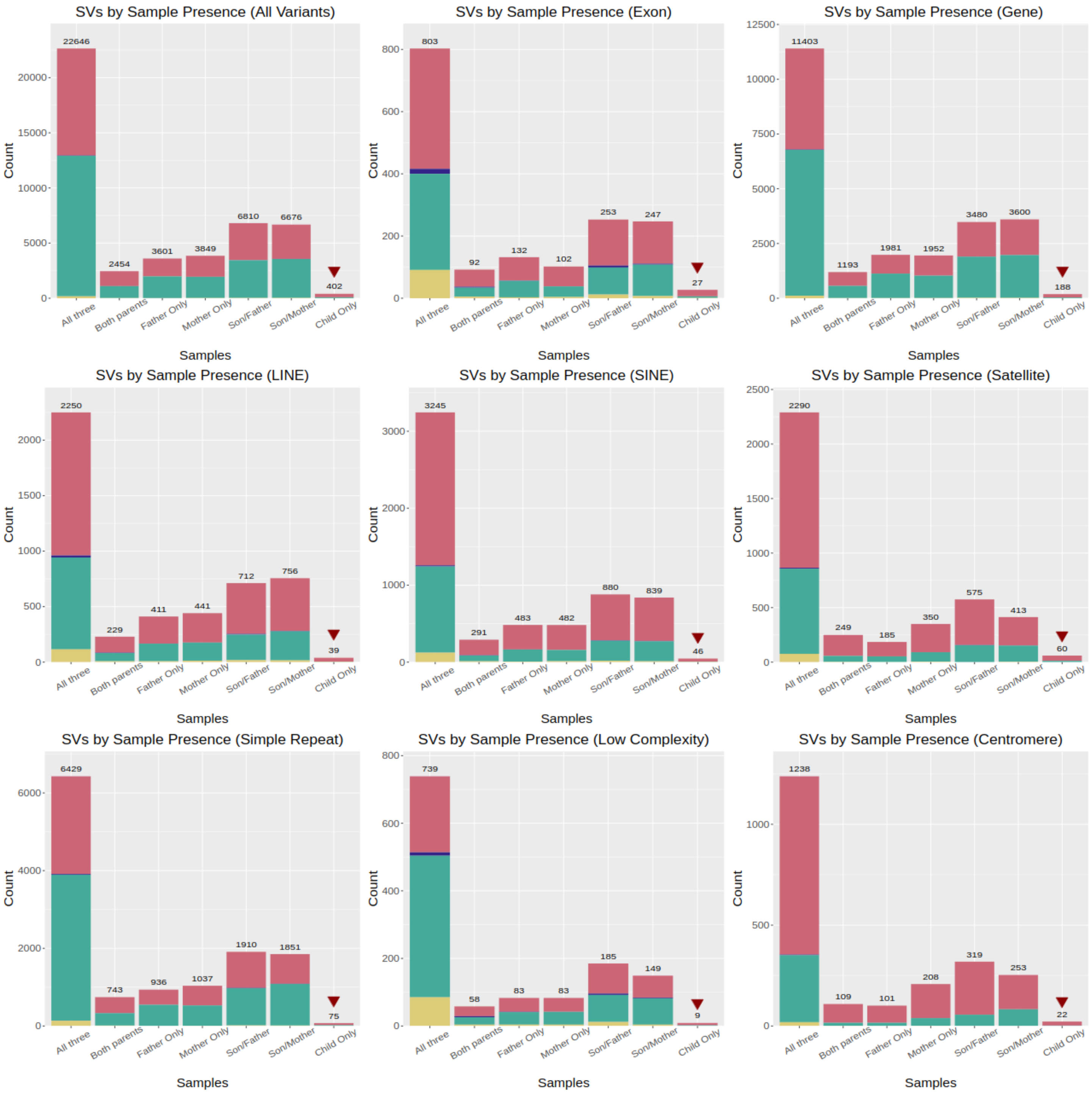


**Supplementary Figure 12. HG002 HiFi Trio Sample Presence by Genomic Context.**

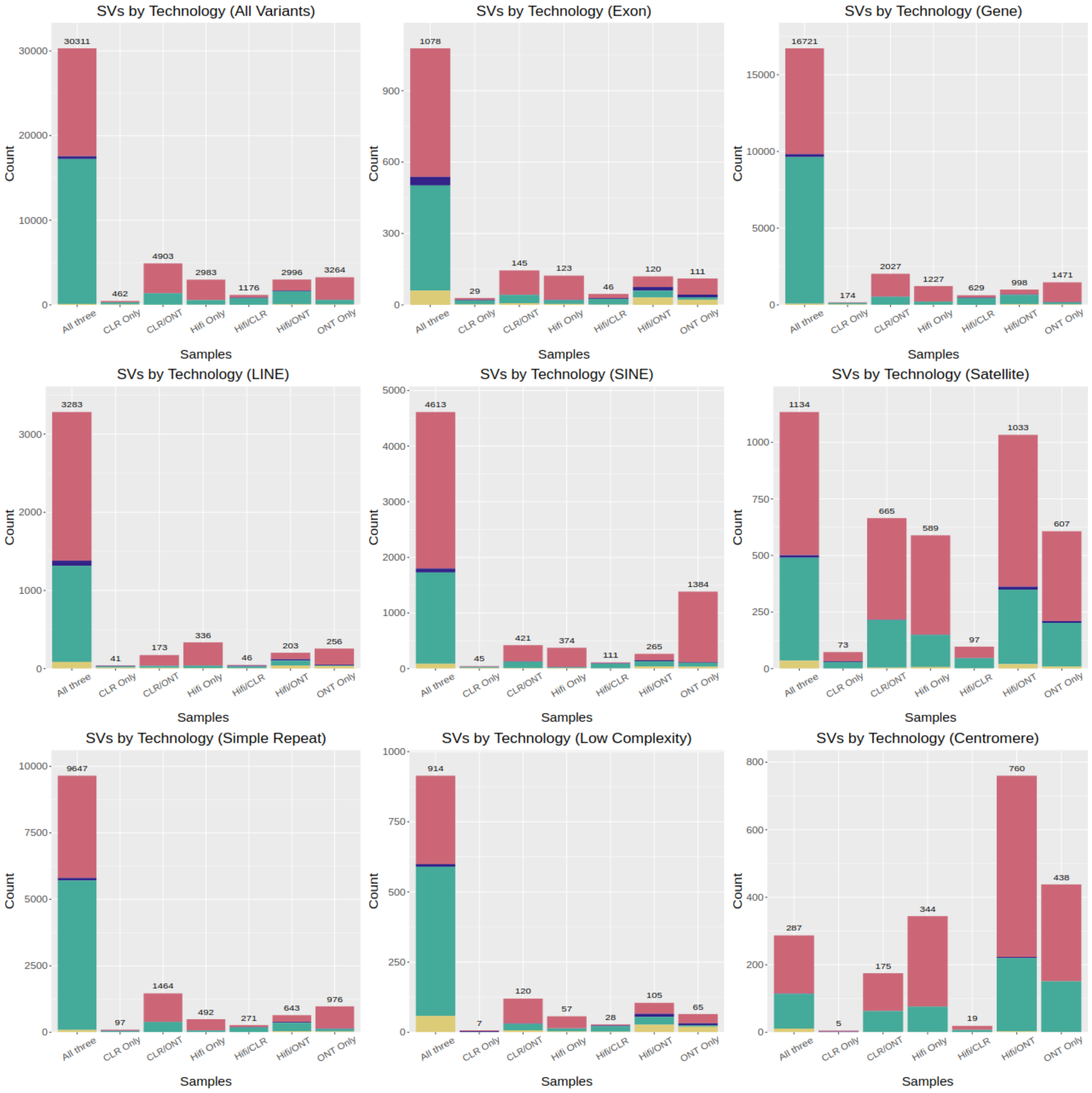


**Supplementary Figure 13. HG002 HiFi Trio Cross-Technology Agreement by Genomic Context.**

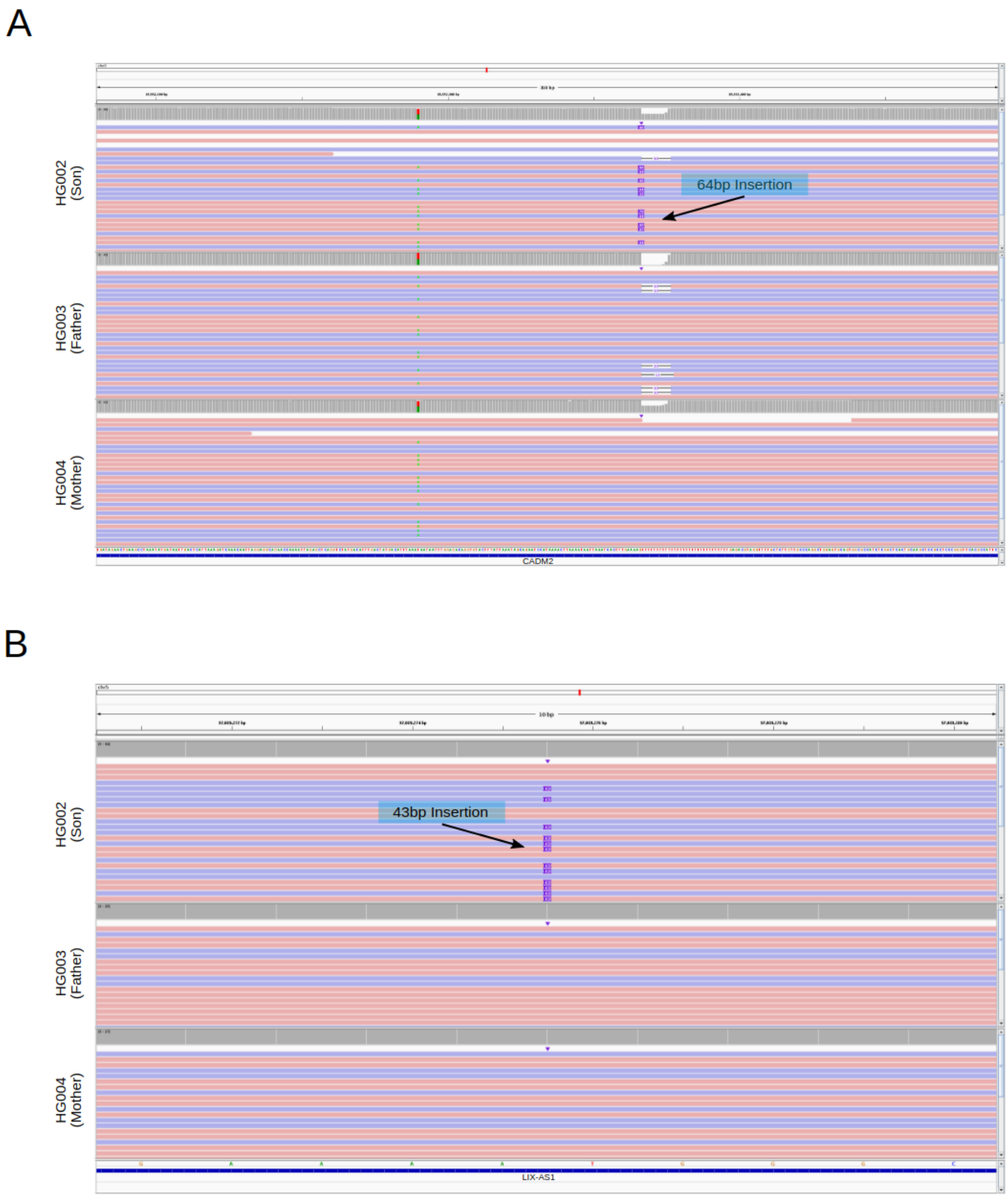


**Supplementary Figure 14. Additional Potential de novo SVs in HG002.**
**a.)** A 64bp insertion at chr3:85552367. This variant was supported as being present in HG002 by HiFi and CLR reads, but missed by the ONT-based calls, likely due to the adjacent homopolymer.
**b.)** A 43bp insertion at chr5:97089276 supported by all three technologies.

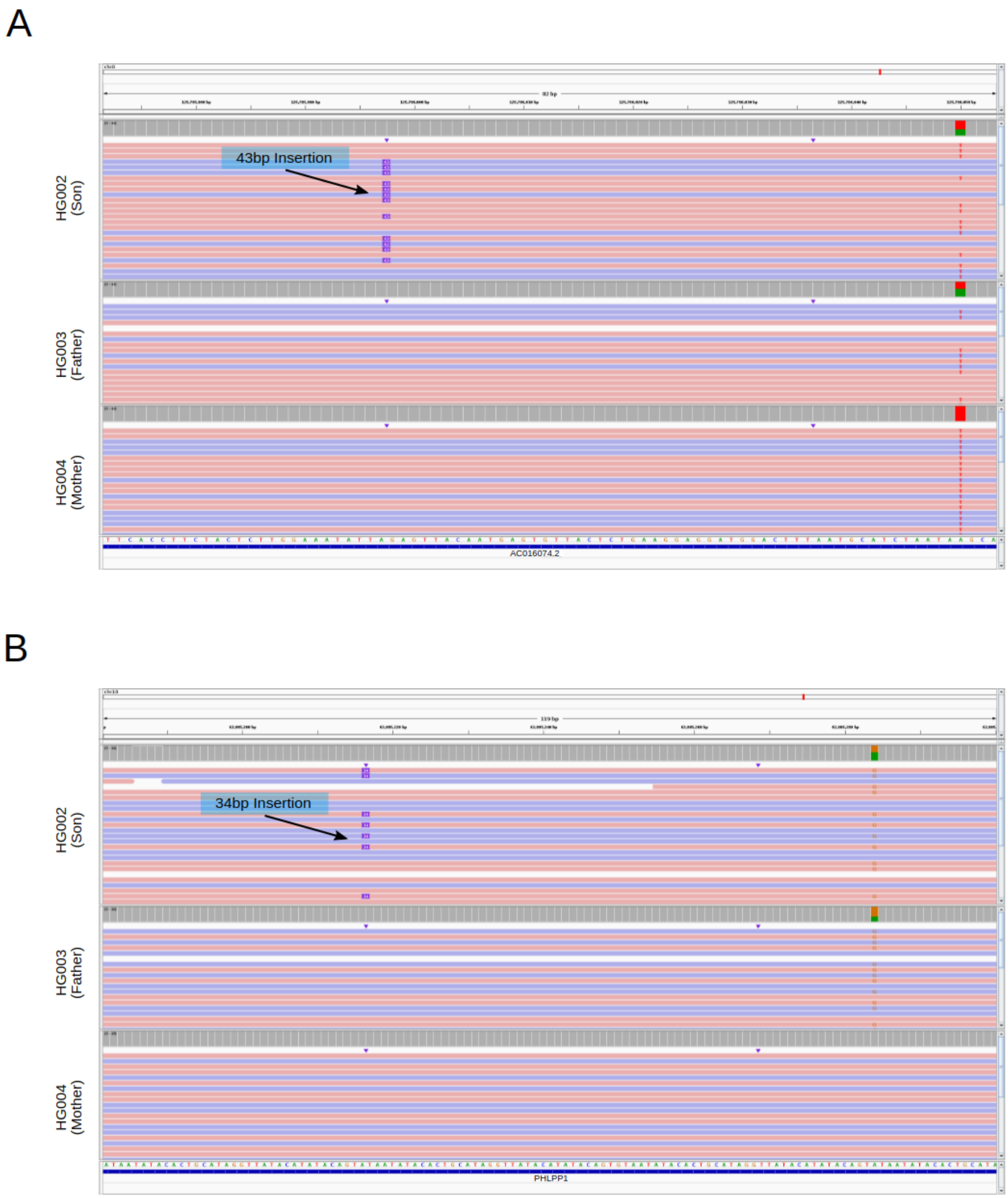


**Supplementary Figure 15. Additional Potential de novo SVs in HG002.**

**a.)** A 43bp insertion at chr8:125785998 on the paternal haplotype supported by all three technologies.

**b.)** A 34bp insertion at chr18:62805217 on the paternal haplotype. This variant was supported as being present in HG002 by HiFi and ONT reads, but missed by the CLR-based calls.

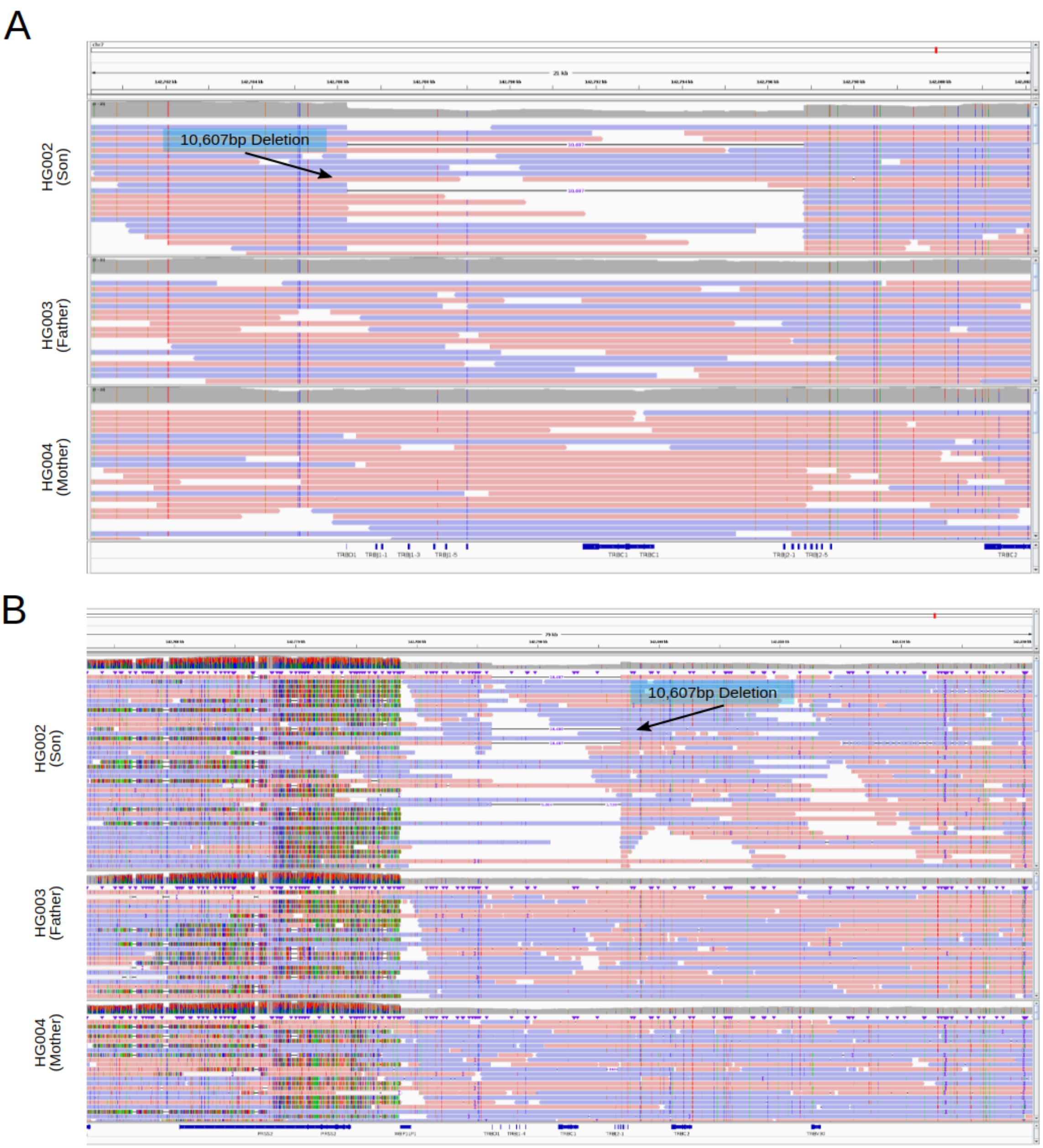


**Supplementary Figure 16. Lower-Confidence Potential de novo SV in HG002.**
This SV is supported by all three technologies as being present in HG002, and is a 10,607bp deletion at chr7:142786222, in the highly variable T cell Receptor Beta (TRB) region. **a.)** IGV screenshot showing the immediate context of the variant. **b.)** IGV screenshot which shows the highly variable region near the variant, leading us to be less confident of the variant being *de novo*.

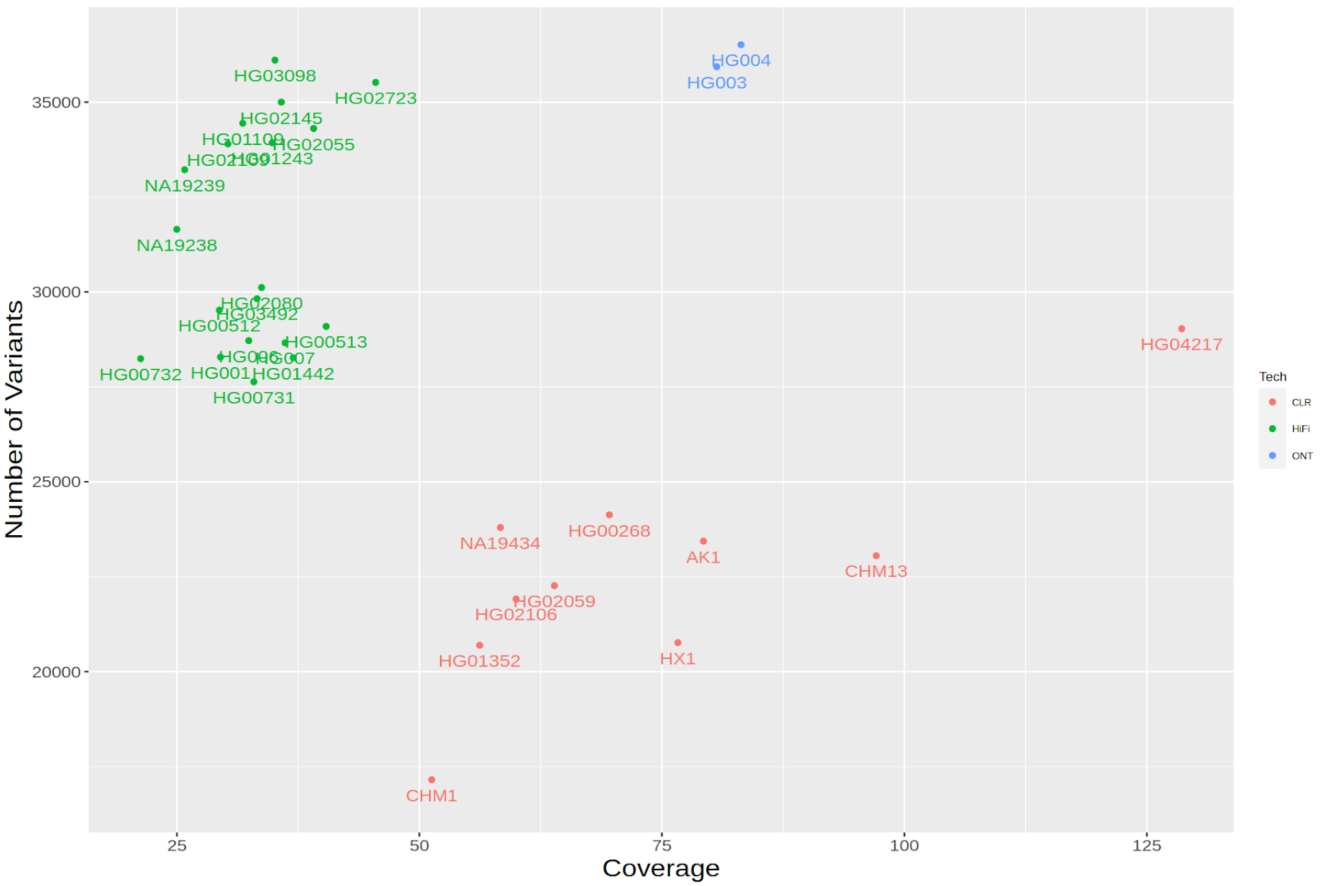


**Supplementary Figure 17. Variant Counts per Sample.**

This shows the number of high-confidence variants called in each sample. While most samples sequenced from the same technology have similar numbers of variants, the coverage of a sample, particularly in those sequenced with CLR, is positively associated with the number of SVs detected.

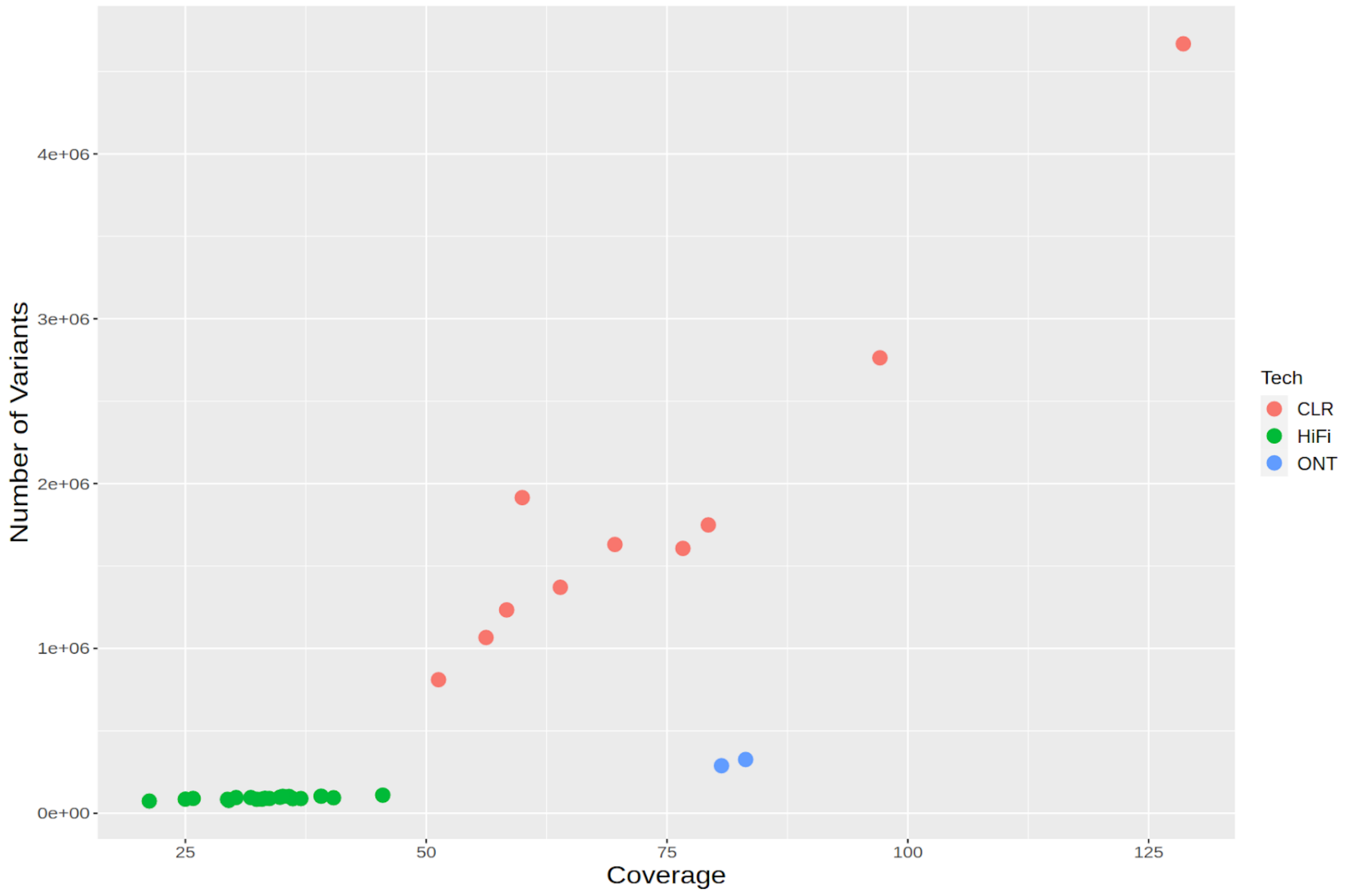


**Supplementary Figure 18. Variant Counts per Sample including Low-Confidence SVs.**
This shows the number of total variants called in each sample with a highly sensitive threshold. There is a high enrichment of low-confidence calls in samples sequenced with CLR, especially samples with high coverage, due to the technology's higher error rate.

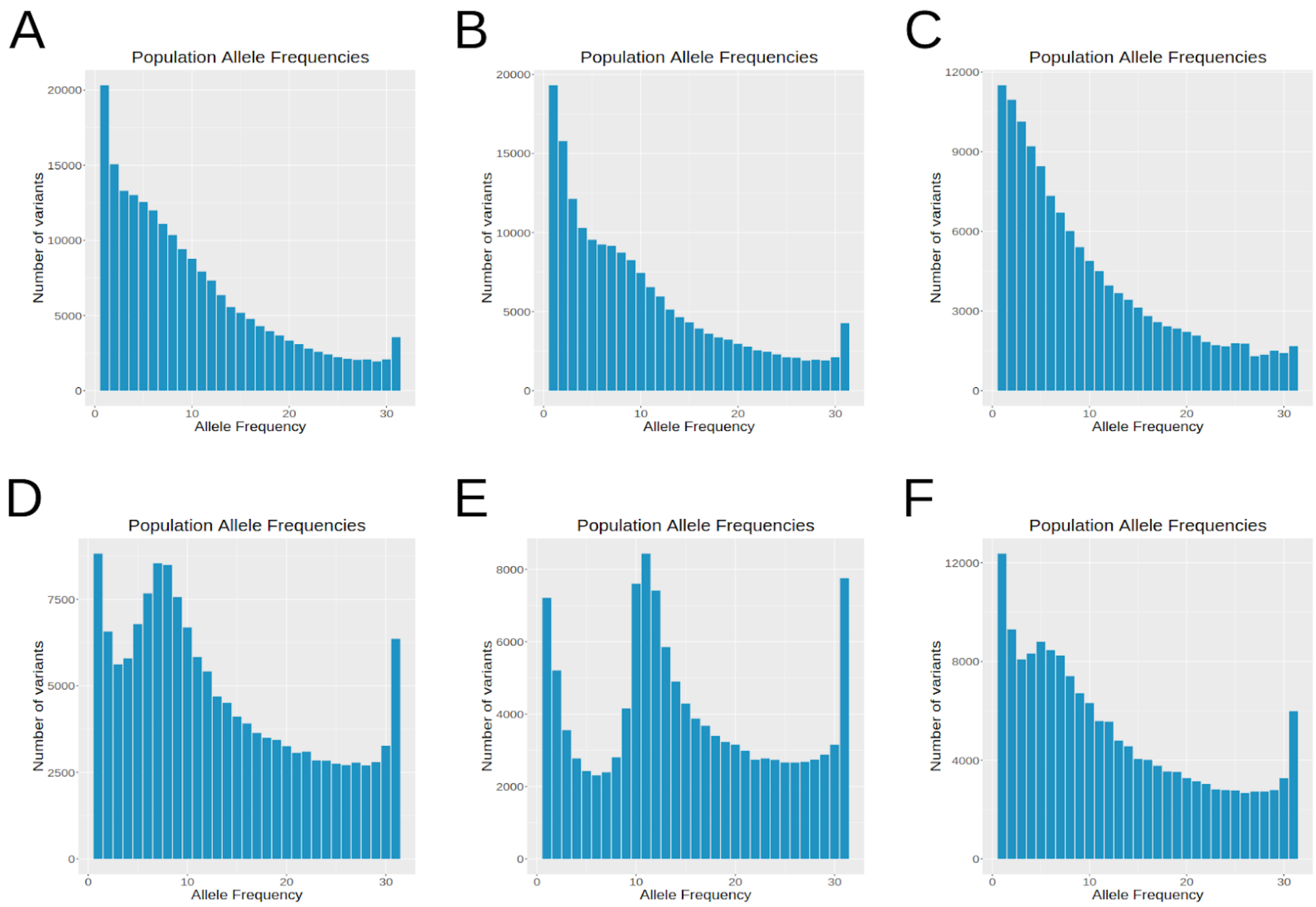


**Supplementary Figure 19. Allele Frequencies of all Merging Software.**
The allele frequency distribution of SVs in the 31-sample cohort when using different methods for merging calls across samples: **a.)** Jasmine **b.)** dbsvmerge **c.)** svpop **d.)** SURVIVOR **e.)** svimmer **f.)** svtools. When using methods which use a constant distance threshold for merging (SURVIVOR, svimmer, svtools), we observe a spike in the allele frequency distribution near 10 samples, where false positive calls from CLR-sequenced samples are merged with each other and with high-confidence variants in other samples.

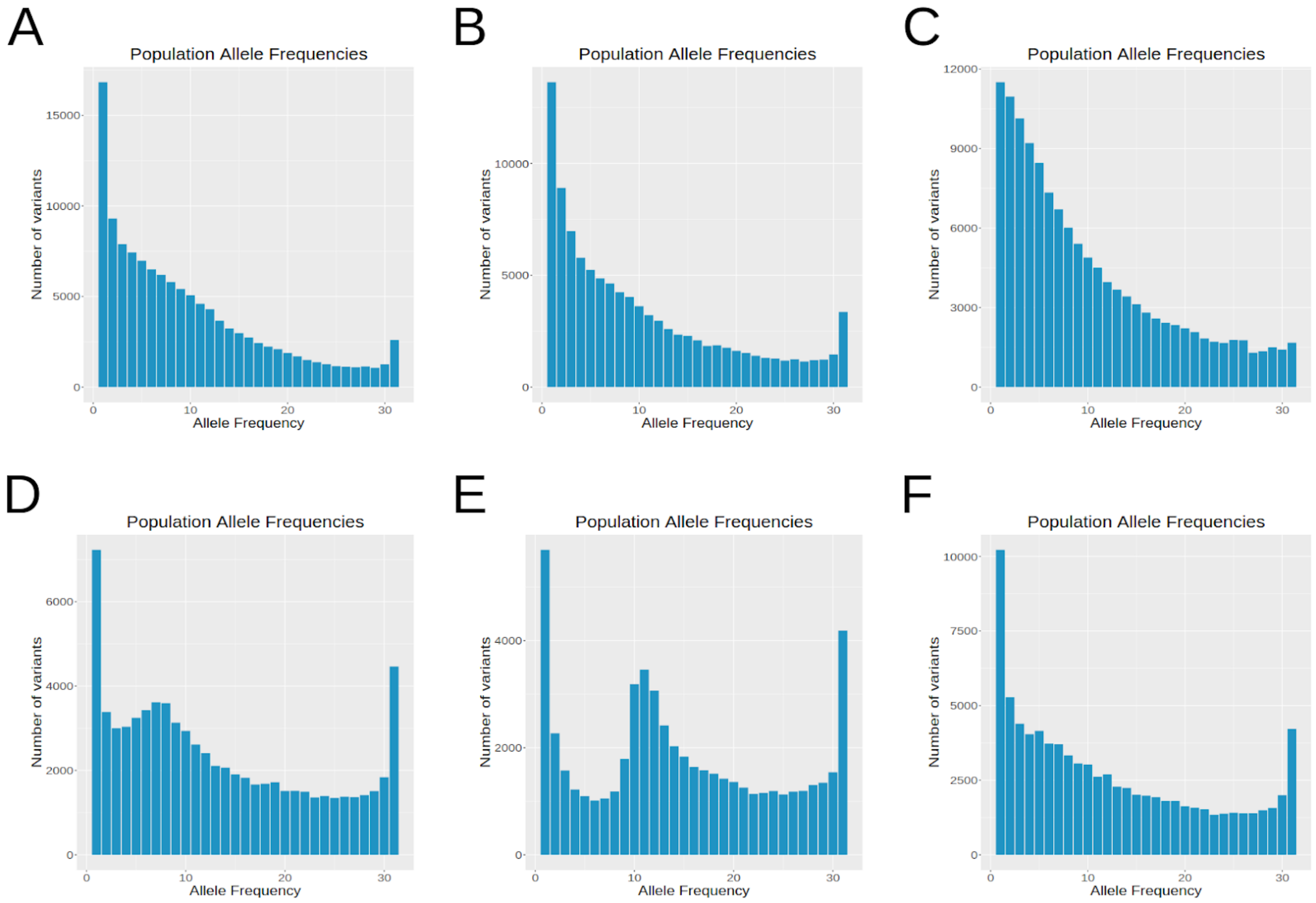


**Supplementary Figure 20. Length-Filtered Allele Frequencies of all Merging Software.**
The allele frequency distribution of SVs in the 31-sample cohort when using different methods for merging calls across samples, filtering out SVs with lengths below 50bp: **a.)** Jasmine **b.)** dbsvmerge **c.)** svpop **d.)** SURVIVOR **e.)** svimmer **f.)** svtools.

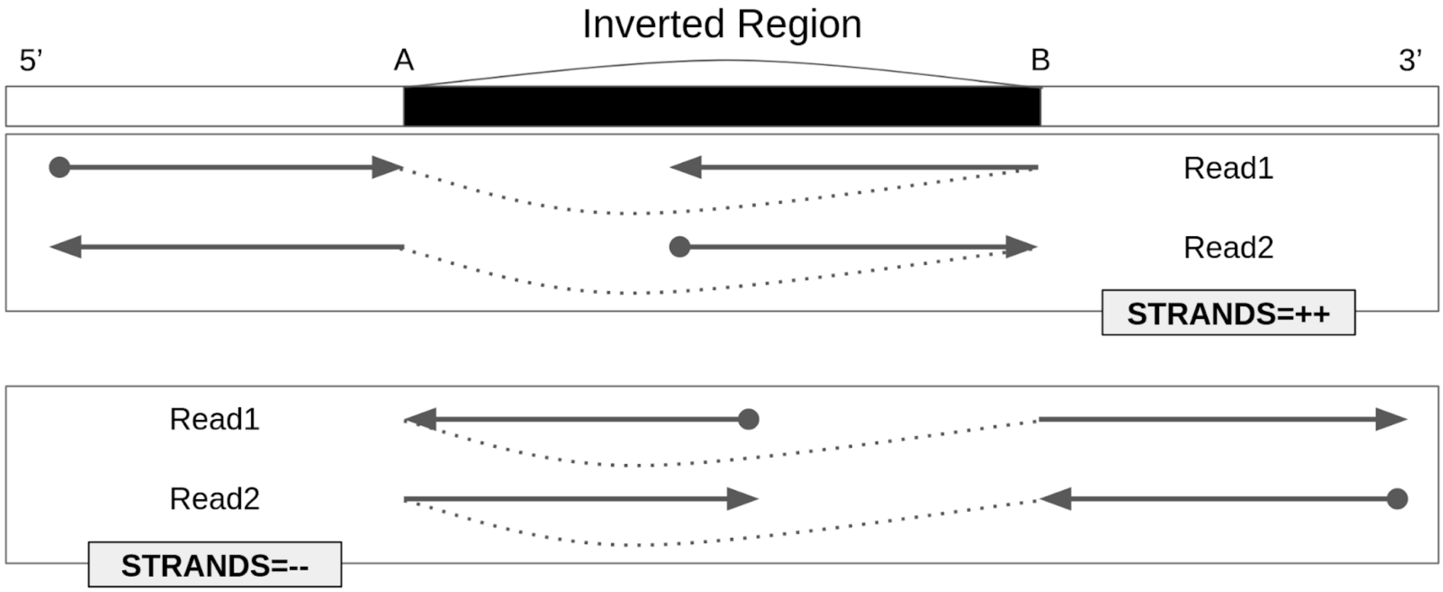


**Supplementary Figure 21. Example of Breakpoint Directionality.**
In complex regions with nested SVs, some variant types such as inversions and translocations, which typically correspond to two novel breakpoint adjacencies, may only be partially present due to other variants interacting with that one. This example shows an inversion where there are two novel breakpoint adjacencies; 1) the sequence near A going towards the 5’ end being newly adjacent to the sequence near B going towards the 5’ end, and 2) the sequence near A going towards the 3’ end being newly adjacent to the sequence near B going towards the 3’ end. While these novel adjacencies typically co-occur, it is necessary to distinguish which is present in the case where only one occurs, so they are reported by sniffles as distinct SV calls with different values for the STRANDS INFO field, and downstream SV comparison/merging software must be aware of the difference between them when comparing across samples.

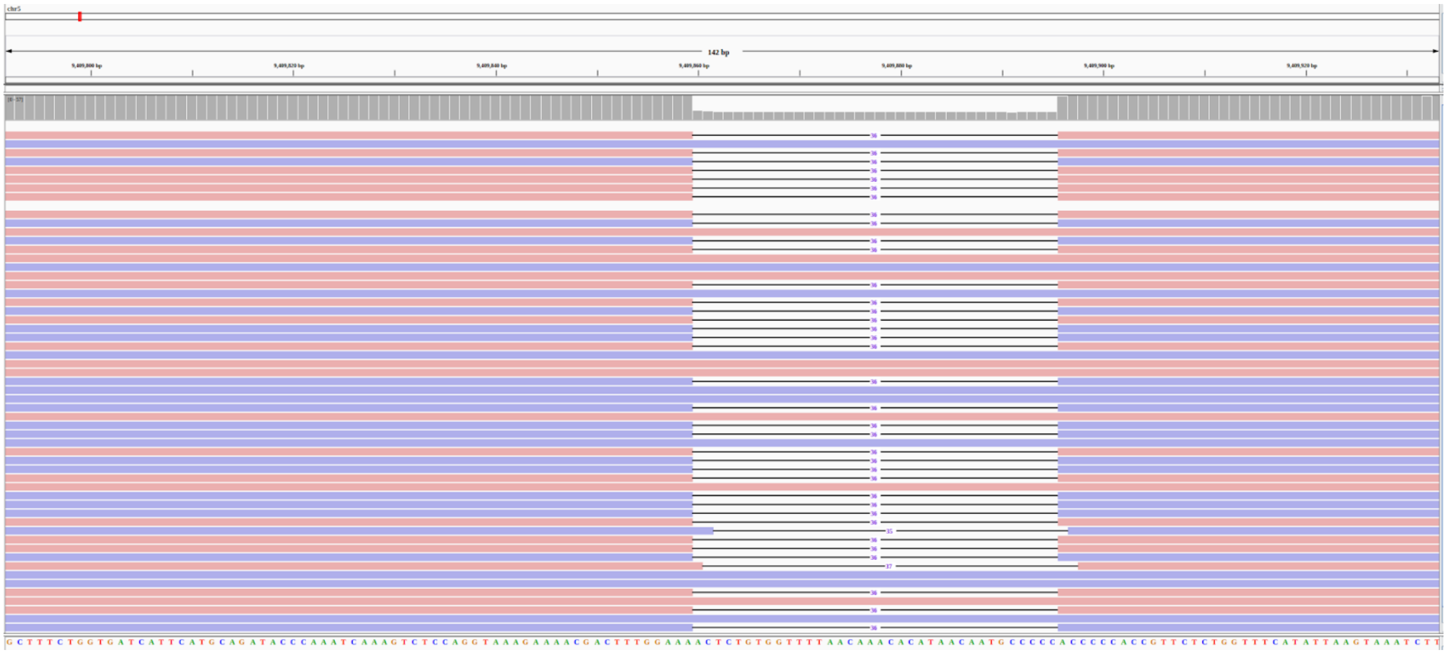

**Supplementary Figure 22. HiFi Alignments for Detecting SEMA5A-Associated SV.**
This shows alignments of HiFi reads in HG01442 at chr5:9409860, the location of the deletion which is associated with the expression of SEMA5A based on the genotypes in 444 samples from the 1000 Genomes Project. HG01442 is one of the nine long-read samples from which the variant was originally called.

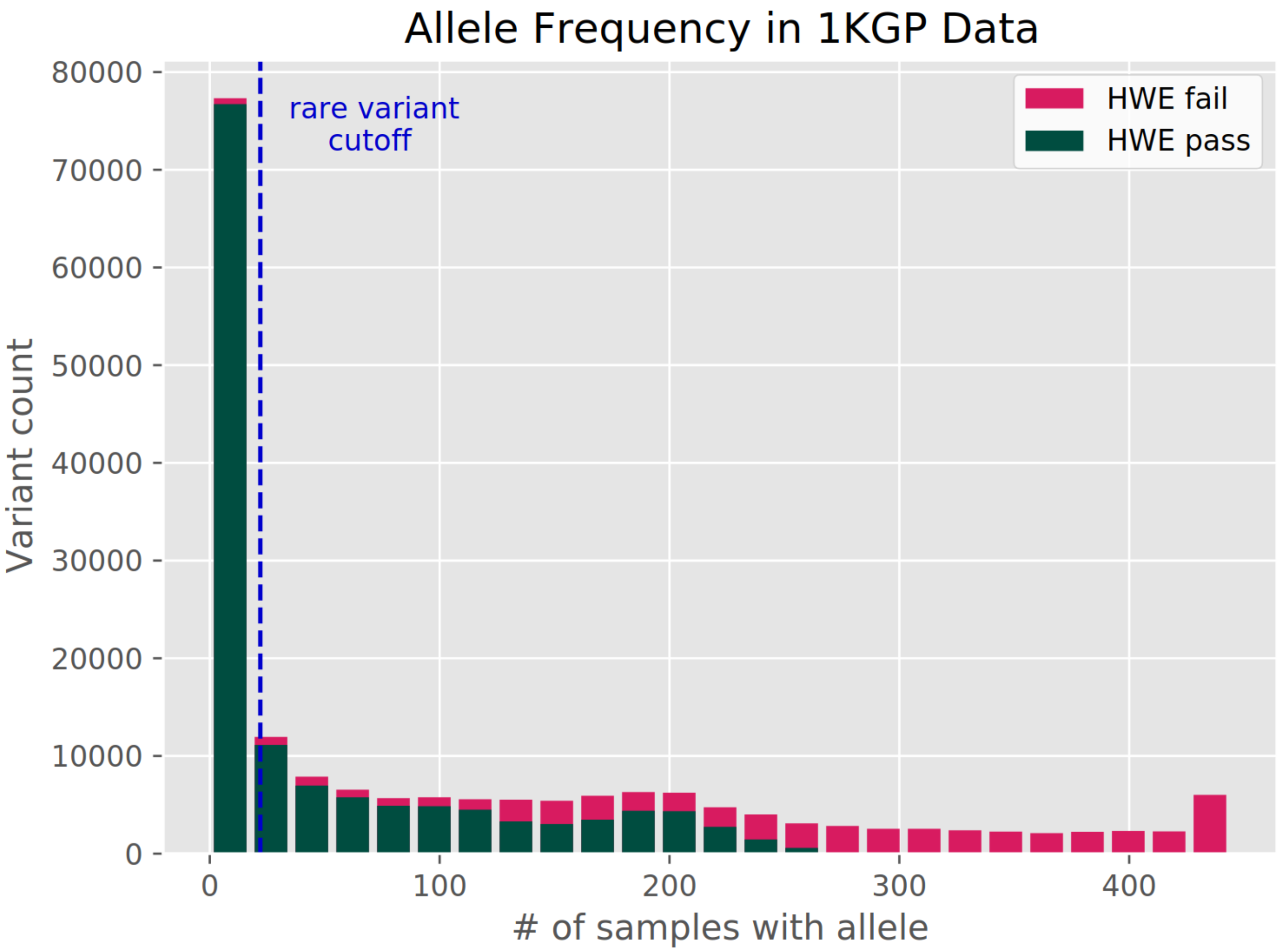


**Supplementary Figure 23. 1KGP Allele Frequencies before and after HWE Filtering.**
Histogram of variant allele frequencies passing and failing the HWE test to filter out variants genotyped by Paragraph in the 1000 Genomes samples that do not match expected Hardy-Weinberg allele frequencies, following best practices (Chen et al. 2019). We later filter out rare variants with allele frequency less than 0.05 (dashed blue line).

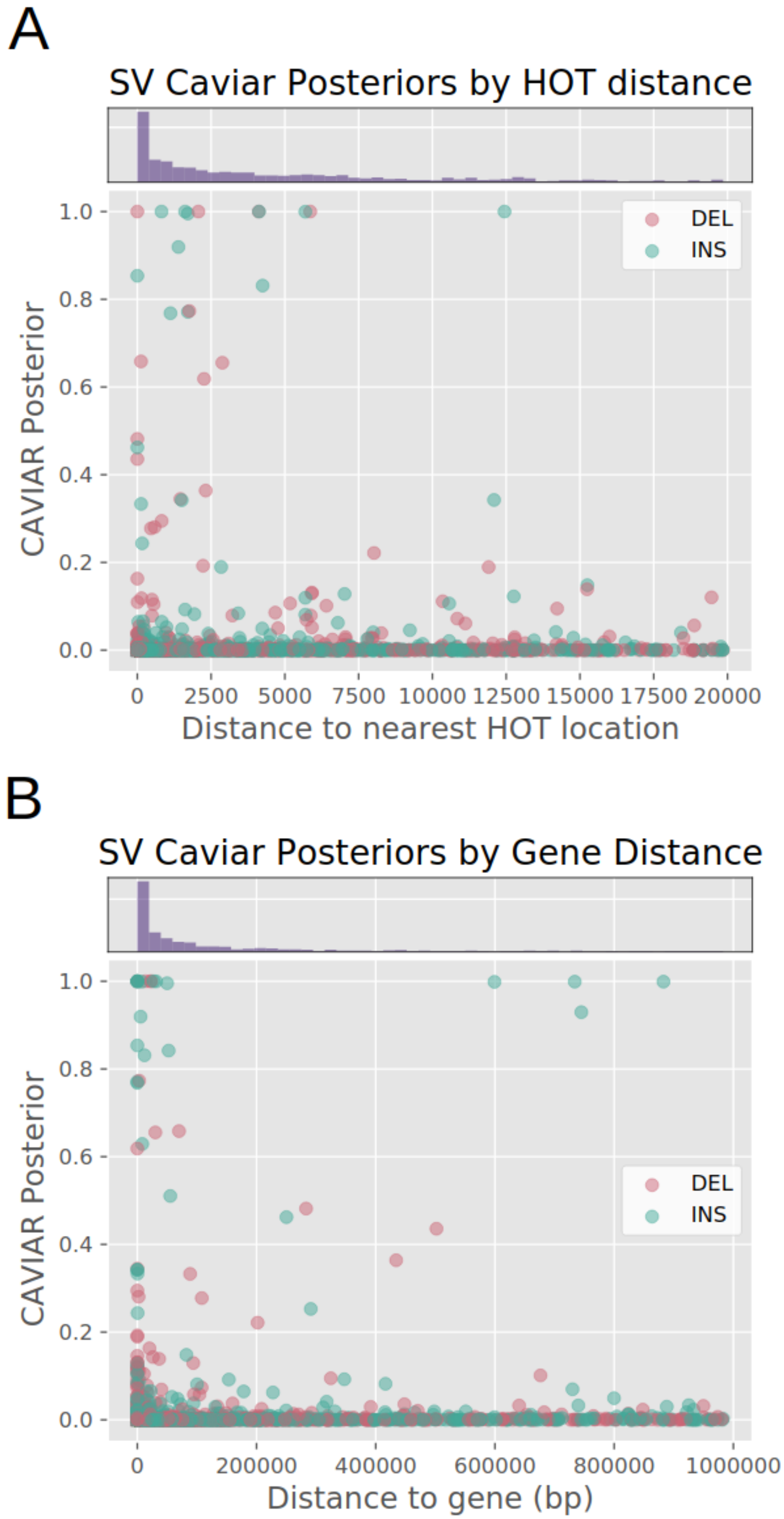


**Supplementary Figure 24. SV Caviar Posteriors by HOT Region and Gene Distance.**
**a.)** Jasmine SV distance to nearest HOT (Highly Occupied by Transcription factors) regions from FunSeq2. Histogram shows distribution of distances to HOT regions. **b.)** Jasmine SV distance to nearest gene. Histogram shows distribution of distances to genes.

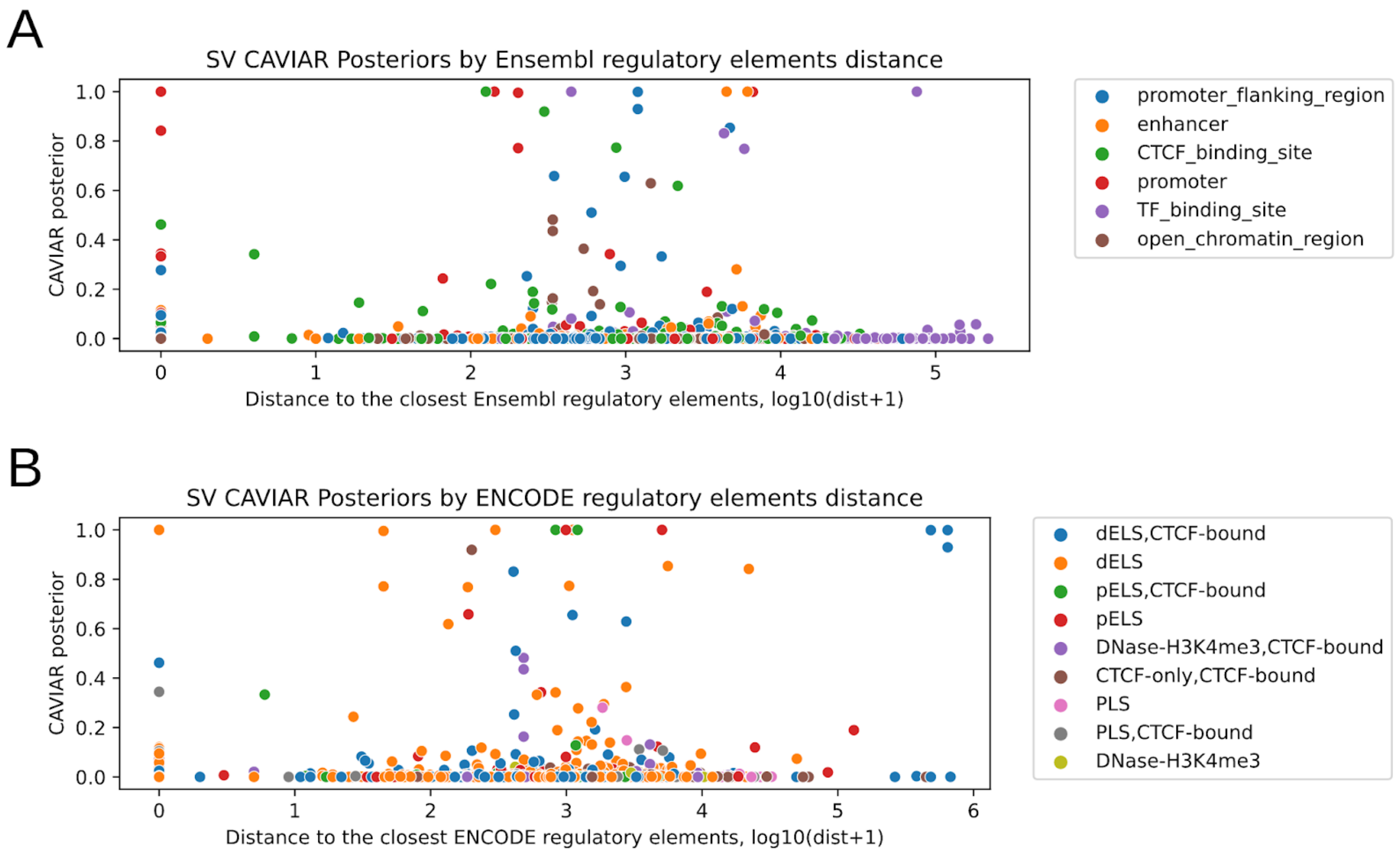


**Supplementary Figure 25. SV CAVIAR Score vs. Distance to Regulatory Elements.** Distances to nearest regulatory elements colored by the type of the nearest elements. **a.)** Log-scaled distances of SVs to various regulatory elements in Ensembl regulatory build. **b.)** Log-scaled distances of SVs to the nearest regulatory elements in ENCODE cis regulatory regions. Both databases are independently derived and most of the SVs with high CAVIAR posteriors are within the 6kb proximal region of regulatory elements. The types of elements near those high-posterior SVs, are primarily promoters, enhancers which are relevant to the regulation of gene expression.

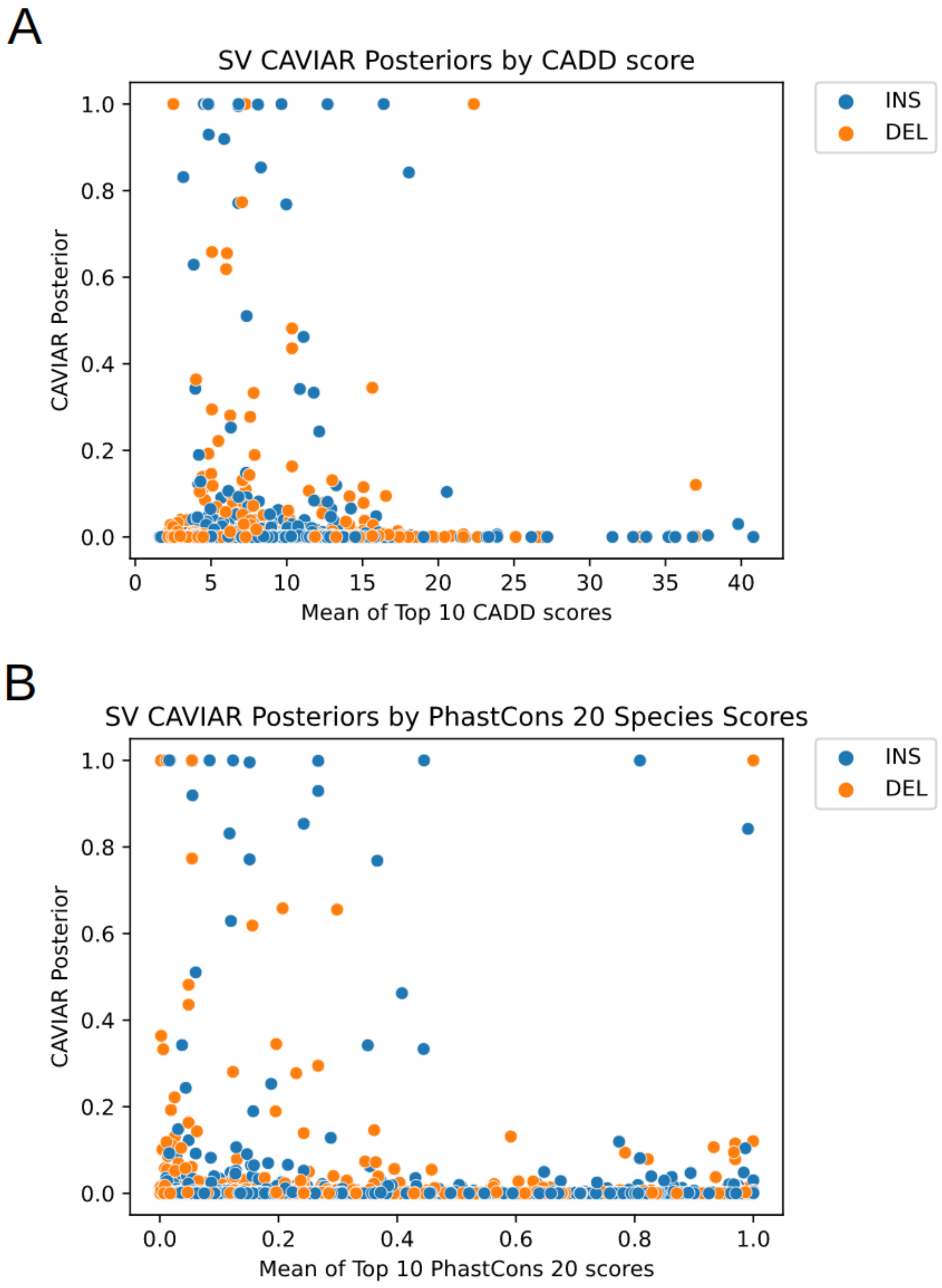


**Supplementary Figure 26. SV CAVIAR Score vs. CADD and PhastCons Scores.**
The mean of the top 10 single-base **a.)** CADD scores scaled as positive Phred-like values and **b.)** PhastCons scores among 20 core mammalian species are calculated in each SV interval. Higher CADD scores indicate higher pathogenic likelihood, while higher PhastCons scores indicate strongly conserved regions. We do not observe evidence of enrichment for conservedness or pathogenicity among SVs with high CAVIAR posteriors.

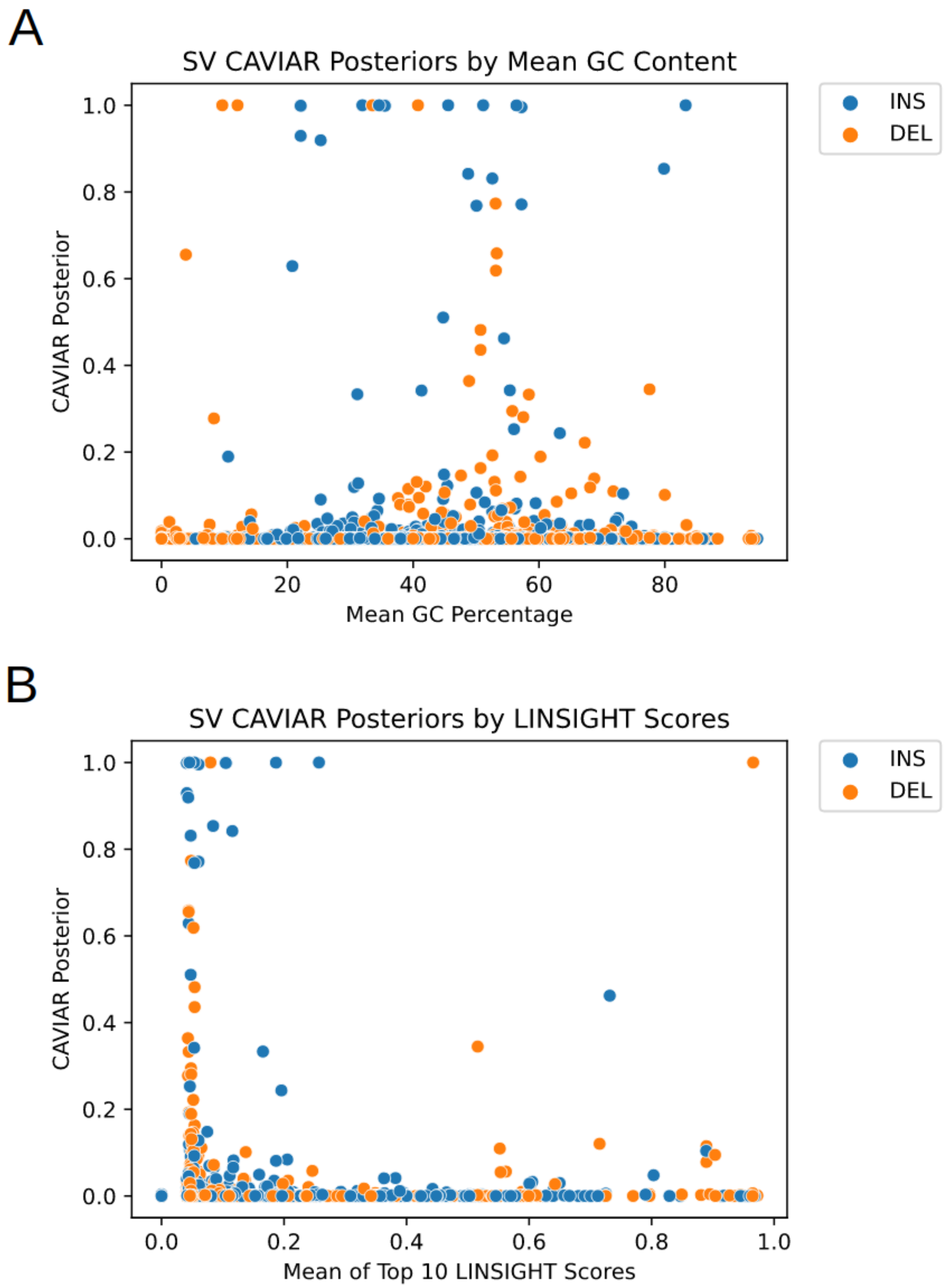


**Supplementary Figure 27. SV CAVIAR Score vs. GC Content and LINSIGHT Score.**
**a)** The mean GC content and **b)** the mean of the top 10 single-base LINSIGHT scores are calculated for each SV interval. GC content is measured as a percentage and is taken from the corresponding track of the UCSC Genome Browser. LINSIGHT scores represent posterior probabilities that each variant has non-coding consequences. We do not observe evidence for SVs with high CAVIAR posteriors to be enriched for extreme GC content or high LINSIGHT scores.

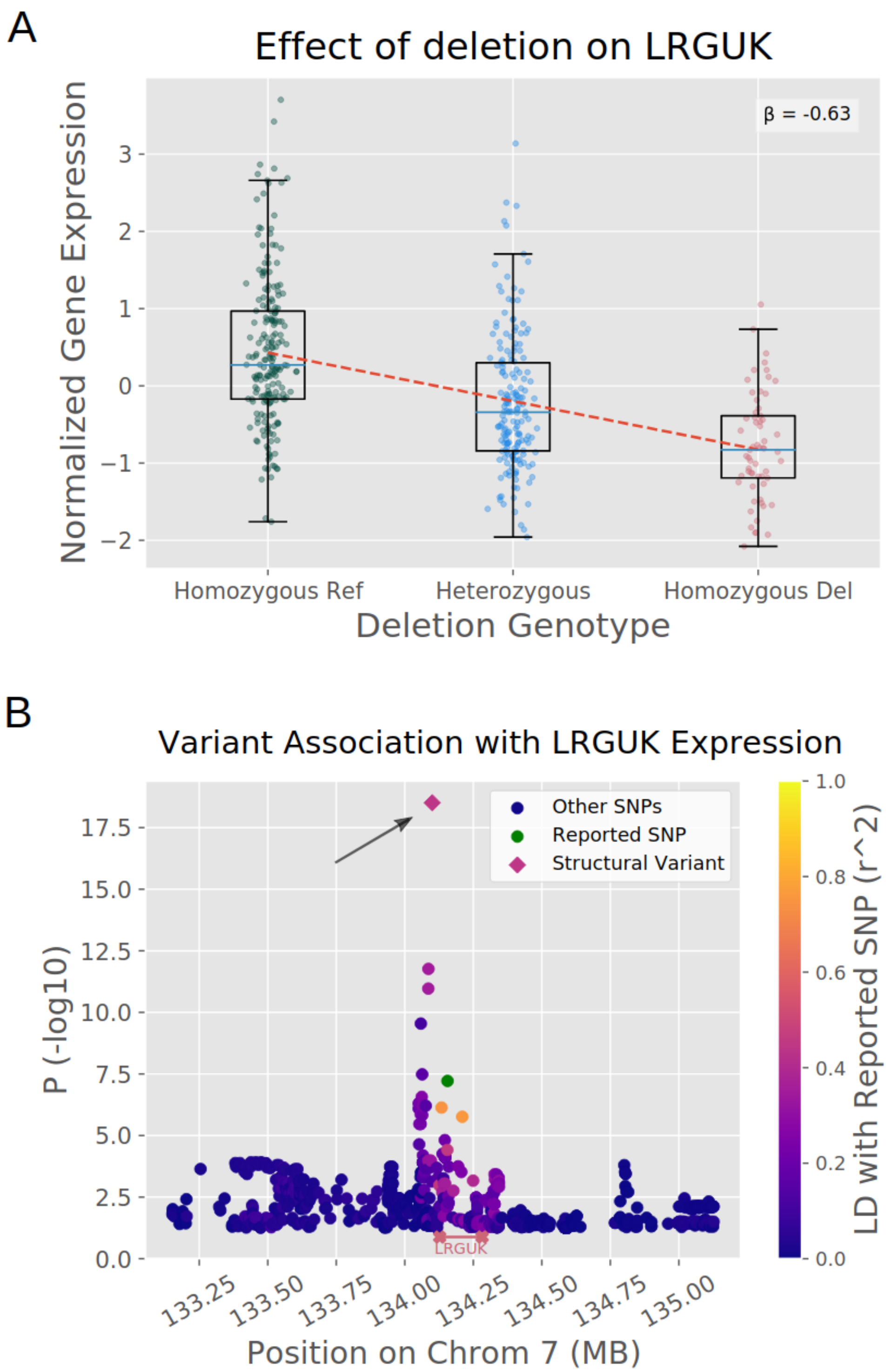


**Supplementary Figure 28. Potential Functionally Relevant Deletion in LRGUK.**
Deletion in LRGUK in LD with reported GWAS SNP. **a.)** Genotype versus gene expression among the 1000 Genomes Project samples for the deletion in LRGUK. **b.)** Manhattan plot for SNPs and novel SV near LRGUK, with p value measured by Wilcoxon rank-sum test. The green point is the SNP reported in GWAS to be associated with smoking initiation (rs1561112), and other points are colored by LD to that SNP.

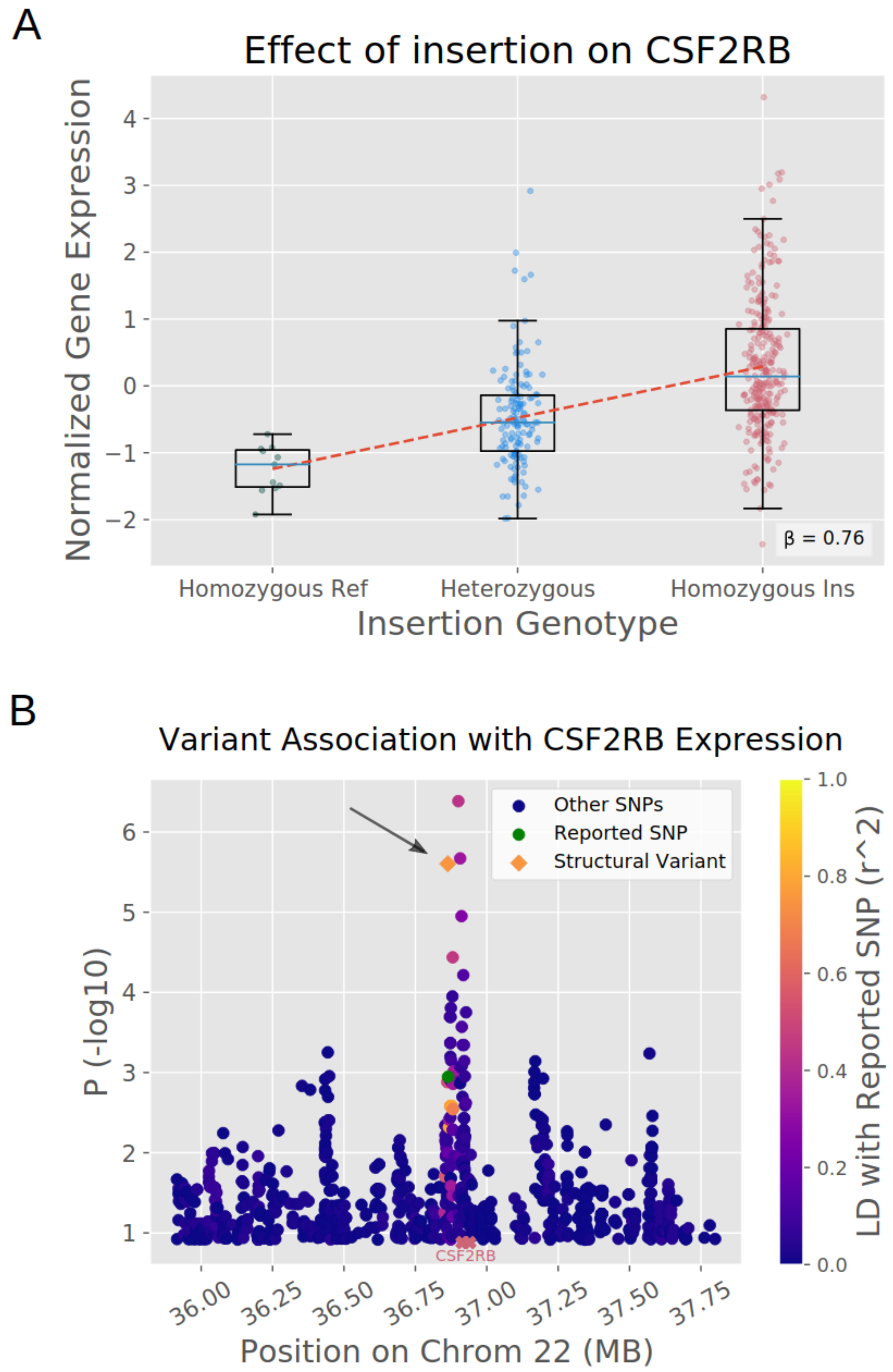


**Supplementary Figure 29. Potential Functionally Relevant Insertion in CSF2RB.**
Insertion in CSF2RB in LD with reported SNP-eQTLs. **a.)** Genotype versus gene expression among the 1000 Genomes Project samples for the insertion in CSF2RB. **b.)** Manhattan plot for SNPs and novel SV near CSF2RB, with p value measured by Wilcoxon rank-sum test. The green point is the SNP-eQTL from GTEx (chr5_9431336_A_T), and other points are colored by LD to that SNP.

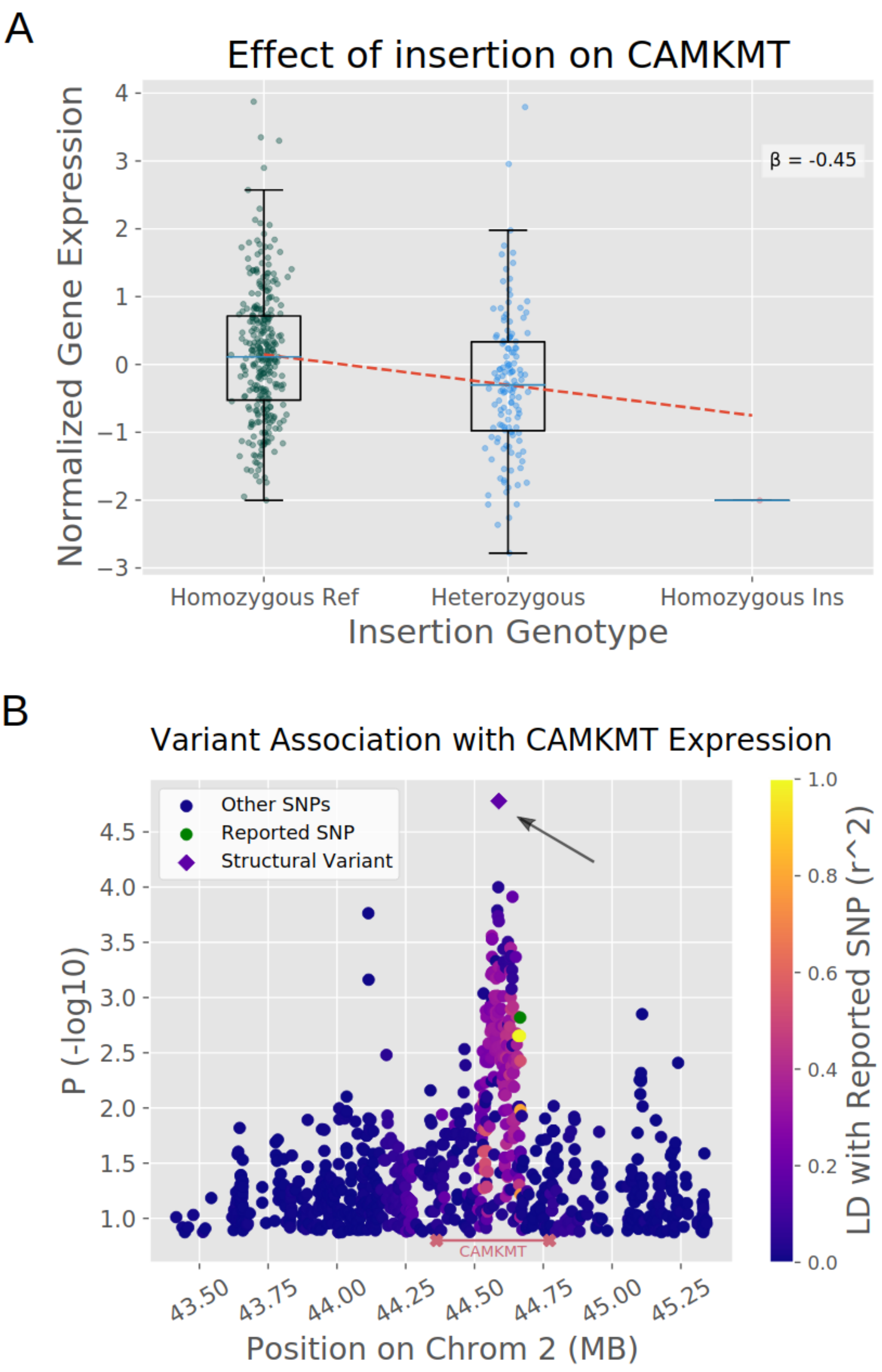


**Supplementary Figure 30. Potential Functionally Relevant Insertion in CAMKMT.**
Insertion in CAMKMT in LD with reported SNP-eQTLs. **a.)** Genotype versus gene expression among the 1000 Genomes Project samples for the insertion in CAMKMT. **b.)** Manhattan plot for SNPs and novel SV near CAMKMT, with p value measured by Wilcoxon rank-sum test. The green point is the SNP-eQTL from GTEx (chr2_44665995_C_T), and other points are colored by LD to that SNP.

### Supplemental Note 1. Jasmine Merging Algorithm.

```
// Takes a set of variants and merges them
MergeAllVariants(vars)
{
    MergedSet = {}

    // Separately merge each chromosome and type
    for(chr in unique({v.chr | v ∈ vars})
    {
        for type in unique({v.type | v ∈ vars})
        {
            SetToMerge = {v | [(v ∈ vars) ∧ (v.chr = chr) ∧ (v.type = type)]}
            MergedSet = MergedSet ∪ MergeSingleGraph(SetToMerge)
        }
    }
    return MergedSet
}



// Merges a set of variants that have the same chromosome and type
MergeSingleGraph(vars)
{
    // Convert SVs to 2D points based on their positions and lengths
    for(v in vars)
    {
        x = v.pos
        y = v.length
        if(v.type == "TRA") y = v.pos2
        v.point2D = (x, y)
    }

    // Add all points to a KD Tree which supports rapid k-nearest neighbor queries
    kdtree = new KDTree()
    kdtree.addAllPoints({v.point2D | v ∈ vars)

    // Initialize a heap to store variant pairs to consider merging
    PairsToProcess = new MinHeap()

    for(v in vars)
    {
        // Initially all variants are in their own component
        v.ComponentId = v.id

        // Initially store the 4 nearest neighbors for each variant
        v.UpcomingNeighbors = kdtree.kNearestNeighbors(v.point2D, 4)

        // Keep track of how many neighbors of each SV have been considered so we know
        // when to refresh the list of upcoming neighbors
        v.NeighborsChecked = 0

        // Add the nearest neighbor for each point to the heap of pairs to consider
        // with the priority key being the Euclidean distance between them
        DistToNearest = EuclideanDistance(v.point2D,  v.UpcomingNeighbors[0])
        PairsToProcess.add(Pair(v, v.UpcomingNeighbors[0].id), DistToNearest)
    }

    // Iterate over the heap until we have no more pairs to consider
    while(PairsToProcess.size > 0)
    {
        NearestPair = PairsToProcess.getMin()

        first = NearestPair.first
        second = NearestPair.second

        // If their distance is bigger than the first point’s threshold we can stop
        // considering any of the first point’s neighbors since they will all be bigger
        if(NearestPair.dist > first.DistanceThreshold) continue

        CanMerge = true

        SamplesWithFirst = unique({v.Sample | v.ComponentID = first.Component})
        SamplesWithSecond = unique({v.Sample | v.ComponentID = second.Component})
```

```
        // If the SVs come from the same sample or have been merged with anything
        // from the same sample, this merge cannot occur
        if(SamplesWithFirst ∩ SamplesWithSecond ≠ ∅)
        {
            CanMerge = False
        }

        // If the distance is too large, the merge cannot occur
        if(NearestPair.dist > second.DistanceThreshold)
        {
            CanMerge - False
        }


        // Perform merging of this variant pair if the merge is valid
        if(CanMerge)
        {
            Merge(vars, first, second)
        }

        // Now get the next nearest neighbor for the first variant in the pair

        first.NeighborsChecked += 1

        // If we used everything we got from the KDTree query, make another
        // query for twice as many neighbors
        if(first.NeighborsChecked == first.UpcomingNeighbors.size)
        {
            NewSize = 2 * first.UpcomingNeighbors.size
            first.UpcomingNeighbors = kdtree.kNearestNeighbors(first.point2D, NewSize)
        }

        // Get the next neighbor from the list and add the pair to the heap
        NextNeighbor = first.UpcomingNeighbors[first.NeighborsChecked]
        DistToNext = EuclideanDistance(v.point2D,  NextNeighbor)
        PairsToProcess.add(Pair(v, NextNeighbor.id), DistToNext)
    }

    // Group variants by component and return
    Results = new Map()
    for(component in unique(vars.ComponentID))
    {
        Results[component] = {v | [(v ∈ vars) ∧ (v.ComponentID = component)]}
    }

    return Results
}


// Merge a pair of variants together by iterating over the smaller component
// and updating their component IDs to match the other variants'.
Merge(vars, first, second)
{
    FirstComponent = {v | [(v ∈ vars) ∧ (v.ComponentID = first.ComponentID)]}
    SecondComponent = {v | [(v ∈ vars) ∧ (v.ComponentID = second.ComponentID)]}

    if(FirstComponent.size > SecondComponent.size)
    {
        for(v in SecondComponent)
        {
            v.ComponentID = first.ComponentID
        }
    }

    else
    {
        for(v in FirstComponent)
        {
            v.ComponentID = second.ComponentID
        }
    }

}
```

**Supplemental Note 2. Commands Used.**

**Jasmine Merging**
```
$ jasmine file_list=FILELIST out_file=MERGED_VCF max_dist_linear=0.5 min_dist=100
```

**dbsvmerge Merging**
```
$ dbSV_merge -f FILELIST -o MERGED_VCF -l 2.0 -r 0.4
```

**svpop Merging**
```
$ python $SVPOPPATH/merge.py FILELIST SAMPLE_LIST MERGED_VCF
```

**SURVIVOR Merging**
```
$ SURVIVOR merge FILELIST 1000 1 1 1 0 1 MERGED_VCF
```

**svtools Merging**
```
$ svtools lsort -r -f FILELIST OUTPUT_SORTED_VCF
$ svtools lmerge -i SORTED_VCF -f 250 > MERGED_VCF
```

**svimmer Merging**
```
$ python svimmer FILELIST CHROMOSOME_LIST --threads 2 --output MERGED_VCF
--max_distance 1000 --max_size_difference 1000 --ids
```

**Paragraph Genotyping**
```
$ multigrmpy.py -m SAMPLE_MANIFEST -i POPULATION_SV_VCF -M [20*DEPTH] -o
OUTPUTFOLDER -r REFERENCE --threads 24 --scratch-dir SCRATCHFOLDER
```

More details of the preprocessing steps and commands used for different merging methods can be found here: https://github.com/mkirsche/SVMergingMethodComparison.

All code for eQTL analysis of SVs can be found here: https://github.com/gautam-prab/jasmine-sv-eqtls.
